# Pancreatic Cancer Intrinsic PI3Kα Activity accelerates Metastasis and rewires Macrophage Component

**DOI:** 10.1101/2020.09.23.307884

**Authors:** B. Thibault, F. Ramos Delgado, E. Pons-Tostivint, N. Therville, C. Cintas, S. Arcucci, S. Cassant-Sourdy, G. Reyes-Castellanos, M. Tosolini, A.V. Villard, C. Cayron, R. Baer, J. Bertrand-Michel, D. Payen, H. Yan, C. Falcomata, F. Muscari, B. Bournet, JP. Delord, E. Aksoy, A. Carrier, P. Cordelier, D. Saur, C. Basset, J. Guillermet-Guibert

**Author notes:** these authors contributed equally to this work (B. Thibault, F. Ramos Delgado, E. Pons-Tostivint). Corresponding author: Julie Guillermet-Guibert, CRCT UMR1037 INSERM-Université Toulouse 3; 2 avenue Hubert Curien; Oncopole de Toulouse; CS, 53717; 31037 TOULOUSE CEDEX 1 - FRANCE, http://www.crct-inserm.fr/17-j-guillermet-guibert-sigdyn-group-pi3k-isoforms-signalling-cancerogenesis-559.html, http://eupancreas.com/julie-guillermet-guibert.

## Abstract

Pancreatic ductal adenocarcinoma (PDAC) patients frequently suffer from undetected micrometastatic disease. This clinical situation would greatly benefit from additional investigation. Therefore, we set out to identify key signalling events that drive metastatic evolution from the pancreas.

We researched a gene signature that could discriminate localised PDAC from confirmed metastatic PDAC and devised a preclinical protocol using circulating cell-free DNA (cfDNA) as an early biomarker of micro-metastatic disease to validate the identification of key signalling events.

Amongst actionable markers of disease progression, the PI3K pathway and a distinctive PI3Kα activation signature predict PDAC aggressiveness and prognosis. Pharmacological or tumour-restricted genetic PI3Kα-selective inhibition prevented macro-metastatic evolution by inhibiting tumoural cell migratory behaviour independently of genetic alterations. We found that PI3Kα inhibition altered the quantity and the species composition of the lipid second messenger PIP_3_ produced, with selective reduction of C36:2 PI-3,4,5-P_3_. PI3Kα inactivation prevented the accumulation of protumoural CD206-positive macrophages in the tumour-adjacent tissue.

Tumour-cell intrinsic PI3Kα therefore promotes pro-metastatic features that could be pharmacologically targeted to delay macro-metastatic evolution.

**The paper explained:** PROBLEM Pancreatic cancer is one of the most lethal solid cancers characterised by rapid progression after primary tumour detection by imaging. Key signalling events that specifically drives this rapid evolution into macro-metastatic disease are so far poorly understood.

RESULT With two unbiased approaches to patient data analysis, higher PI3K pathway and more specifically higher PI3Kα activation signature can now be identified in the most aggressive pancreatic cancer primary tumours, that lead to earlier patient death. Our in vitro data showed that PI3Kα is a major positive regulator of tumour cell escape from the primary tumour: tumour-intrinsic PI3Kα activity enables actin cytoskeleton remodelling to escape the pancreatic tumour. We chose to use two preclinical models of pancreatic cancer to validate that PI3Kα is a target for delaying evolution of PDAC. The first one mimicked pancreatic patient micrometastatic disease that is undetected by echography and consisted in treating mice presenting echography detected primary tumours combined with increased circulating DNA as a blood biomarker of the most aggressive tumours. The second model consisted in studying the tumour cell implantation and their early proliferation in metastatic organ after injection in blood. We treated both preclinical models with a clinically relevant PI3K α-selective inhibitor (BYL-719/Alpelisib), that is currently being tested in pancreatic cancer patients (without any patient selection). We found that PI3Kα activity drives evolution of micrometastatic disease towards macro-metastatic stage in both models: inhibition of PI3Kα delayed primary tumour and micro-metastasis evolution. Finally, PI3Kα activity increases protumoural characteristics in peritumoural immune cells via tumour cell-intrinsic cytokine production that could facilitate metastatic evolution.

IMPACT Circulating tumour DNA represents a strong independent biomarker linked to relapse and poor survival in solid cancer patients. A clinical study in resected PDAC patients with micrometastatic disease characterised by high circulating tumoural DNA levels is needed to assess if PI3Kα-selective inhibitors significantly delay metastatic progression and death.

**Graphical Abstract:** Pancreatic ductal adenocarcinoma requires tumour-intrinsic PI3Kα activity to accelerate inflammatory metastatic disease.
Biorender illustration.

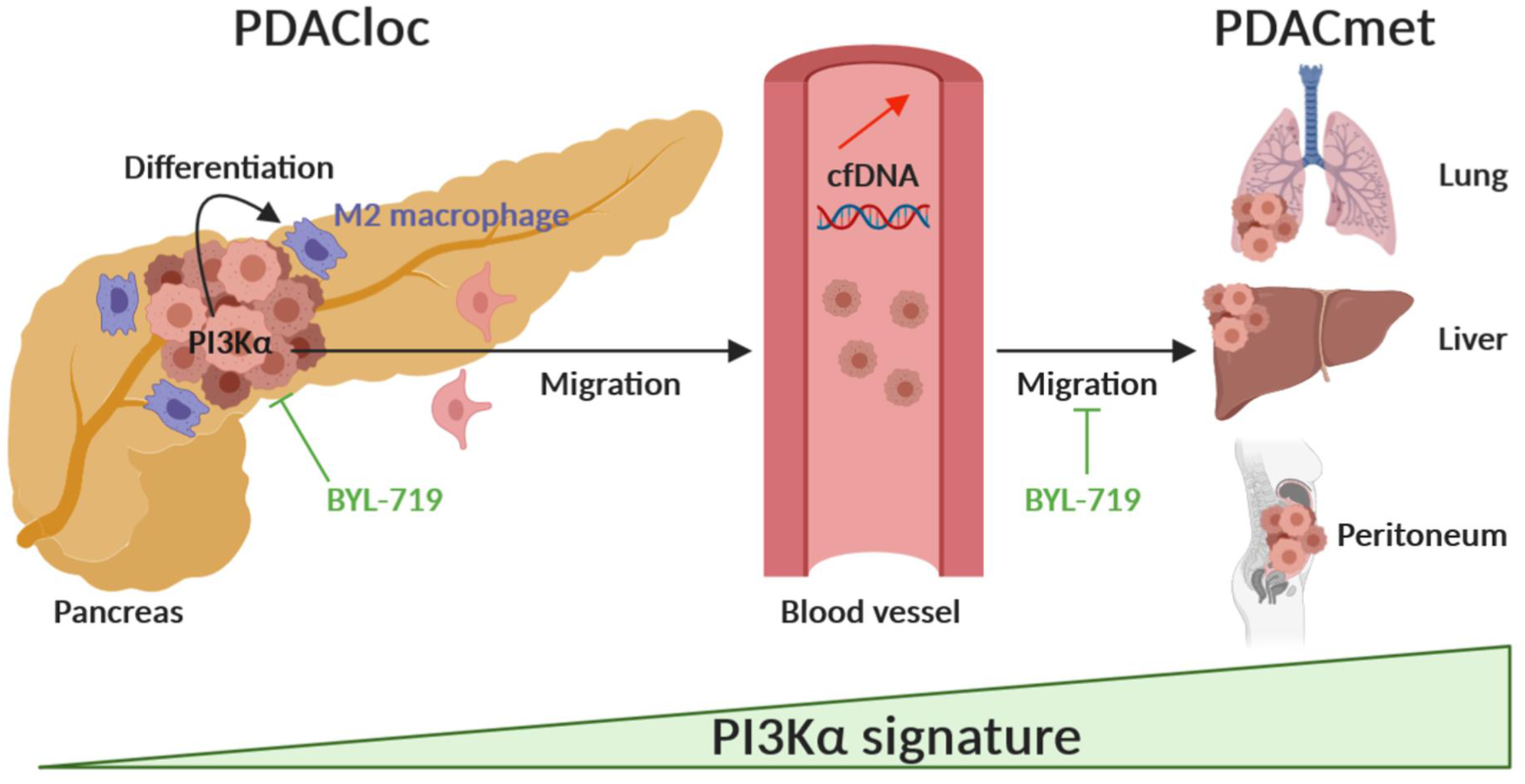

## Introduction

In humans, PI3Ks are composed of 8 isoforms distributed into 3 classes. Each class I PI3K dimers, namely called PI3Kα, PI3Kβ, PI3Kγ and PI3Kδ are composed of a catalytic subunit (p110α, p110β, p110γand p110δ) and a regulatory subunit (p85 for α, β and γ and p101/p87 for γ). PI3Kα and PI3Kβ are ubiquitously expressed. PI3Kγ and PI3Kδ are restricted to the cardiovascular system and leukocytes in normal tissues and can be found overexpressed in solid tumours (1). PI3Ks are lipid kinases that phosphorylate phosphatidyl inositol 4,5 – biphosphate (PIP_2_) into PIP_3_ which acts as a second messenger and regulates various functions in normal and tumour cells *via* the PI3K/Akt/mTOR pathway. The PI3K/Akt axis is frequently hyper-activated in cancers and has been tested as a clinical target in recent years (2, 3). PI3K inhibitors are currently described as cytostatic agents, since PI3K activity is critically driving oncogenesis in a cell-autonomous manner. However, the clinical importance of other cell functions regulated by this pathway in a non-cancer cell autonomous manner, particularly on macrophages, has been underestimated.

Pancreatic ductal adenocarcinoma (PDAC) is a lethal cancer (4) where activation of class I PI3K is high and linked to poor prognosis (5). Localised, locally advanced and metastatic PDAC are characterised by early surgical relapse and failure of long-term disease control with chemotherapies, respectively. Molecular characterisation of large cohorts of PDAC patients demonstrates that oncogenic *KRAS* mutations on G12 position are found in more than 80% of all patients. There are multiple altered signalling pathways downstream of oncogenic KRAS, including PI3K/Akt pathway (6, 7). Fewer than 5% of patients present *PIK3CA* oncogenic mutation, but this mutation mimics the KRAS oncogenic pathway (8). Our research group and others have demonstrated that the lipid kinase PI3Kα drives the initiation of pancreatic cancer downstream of oncogenic KRAS (9, 10). However, little is known of the importance of this PI3K isoform in progression of existing tumours towards metastatic disease.

Cell-free DNA (cfDNA) and, more precisely, circulating tumour DNA (ctDNA) appears in clinical oncology as an attractive biomarker for early cancer detection, diagnosis and prognosis (11–13). In cancer patients, ctDNA represents a variable fraction of cfDNA (12) and is distinguished by the presence of specific cancer-associated mutations. The release of cfDNA can be due to apoptosis and necrosis of cancer cells (or healthy cells), and it can be secreted directly by the tumour or microenvironment cells such as immune and inflammatory cells (15). Since cfDNA has been studied as an exploratory biomarker of micrometastatic disease in PDAC (16, 17), we propose that detection of cfDNA as a sign of early metastatic disease could predict therapeutic efficiency towards metastatic evolution.

With two unbiased approaches to patient data analysis, we demonstrated that gene expression signatures of PI3K activation were a novel way to molecularly identify the most aggressive primary tumours. We then identified a novel pharmacological target, PI3Kα, that drives pro-inflammatory features towards macro-metastatic evolution. PI3Kα inhibitors could be included in the immunomodulatory and anti-cancer therapeutic arsenal in PDAC.

## Results

### PI3K and PI3Kα-specific transcriptomic signature predicts aggressive pancreatic cancer

We sought to determine in an unbiased manner which signalling pathway are associated with aggressive features in PDAC.

We analysed publicly available dataset to distinguish a normal pancreas from chronic pancreatitis (CP) and primary tumours from localised PDAC (PDACloc) or metastatic PDAC (PDACmet) (Fig. 1a-c, suppl. Table 1). PDACmet and CP patients share the same enrichment of mRNA expression-based hallmarks of biological pathways compared to normal, except for 3 hallmarks. The PI3K/Akt/mTOR was the pathway differentially expressed with the lowest p value (Fig. 1a). Differential enrichment analysis demonstrates that the PI3K/Akt/mTOR pathway is constantly (equally) enriched in localized and metastatic PDAC compared to normal parenchyma. This trend was confirmed following Reactome pathway analysis; however, we found that PI3K cascade to FGFR2 was significantly increased in PDACmet as opposed to PDACloc, suggesting differential activation of receptor tyrosine kinase (RTK)-coupled PI3K in these samples. As PI3Kα is key for insulin signalling (18), angiogenesis (19) and PDAC initiation (9), we then designed a PI3Kα activation gene signature, based on expression levels of PI3Kα-regulated curated genes (suppl. Fig. 1). PI3Kα activation scoring allowed us to cluster 8/9 PDACmet patients (Fig. 1c). Conversely, only 2/9 PDACloc clustered with PDACmet. We extended our findings to two larger, independent cohorts of PDAC patients (suppl. Table 2). High scoring of PI3Kα activation was significantly increased in patients with the poorest prognosis, regardless of their stage (Fig. 1d). The PI3K activation signature discriminates between localised patients and those with an early risk of relapse and death (Fig. 1e). Thus, even though oncogenic mutations of PI3Kα are rare in PDAC (8), as confirmed in the PAAD database (suppl. Table 3, suppl Fig. 2), high scoring of non-mutated PI3Kα activation was a worse prognostic factor, irrespective of the stage of the disease. In conclusion, activation of non-mutated PI3Kα appears as a strong prognostic factor of aggressive disease.

**Figure 1:**
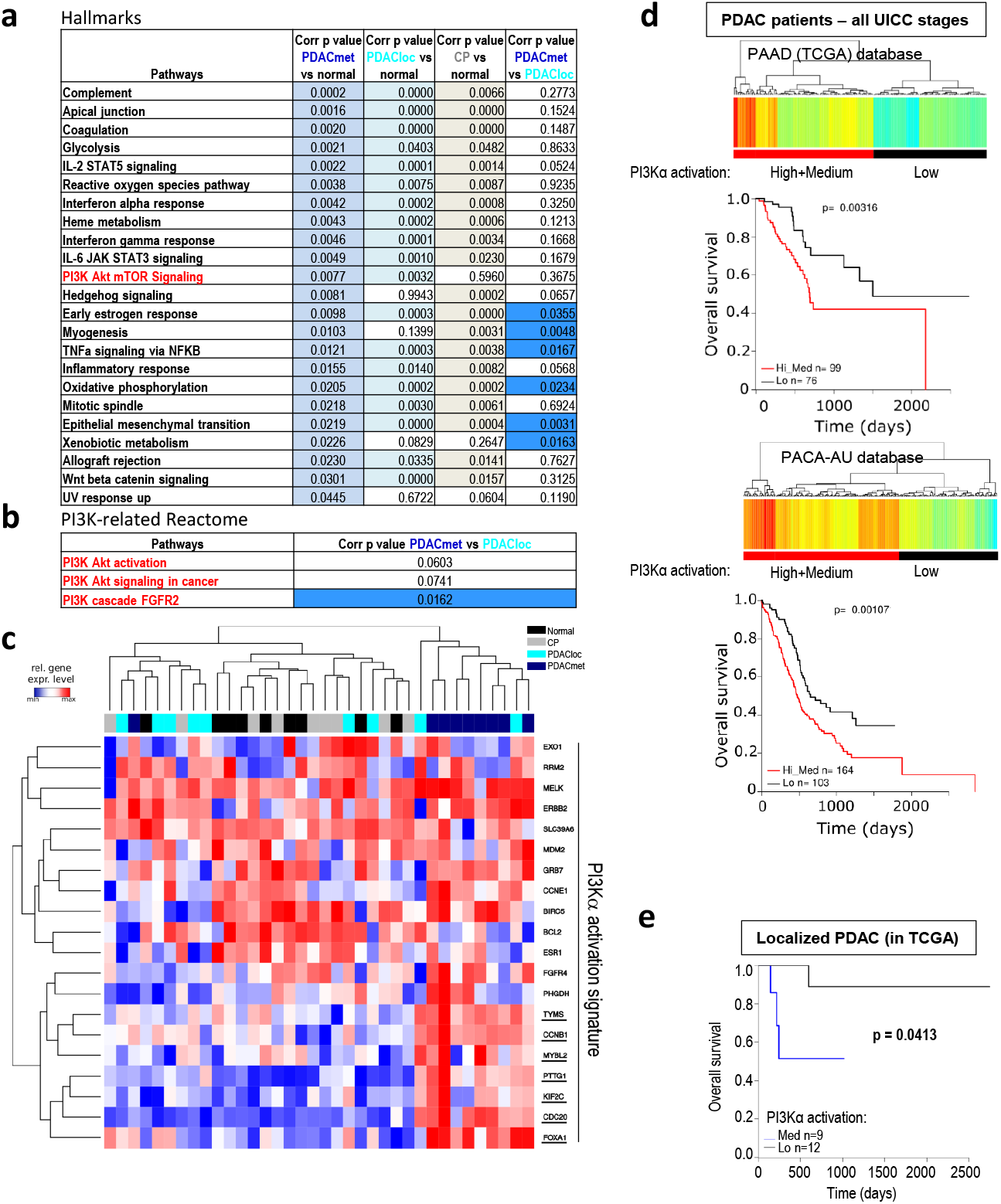
A PI3Kα specific transcriptomic signature predicts pancreatic cancer aggressiveness. **a**, List of 19 HALLMARK pathways (out of 50) significantly altered (ranked with p value – top: low, bottom: high) in a cohort of 9 metastatic pancreatic adenocarcinoma patients (PDACmet), 9 localised PDAC (PDACloc), 8 chronic pancreatitis (CP) compared to 9 normal pancreas samples (normal); corrected p values correspond to the comparison indicated. Significantly altered corrected p values are highlighted in colour. **b**, List of PI3K related pathway gene signatures (Reactome) altered in the same cohorte; corrected p values are shown. **c**, Transcriptomic signature indicative of PI3Kα activation in the same cohorte. Genes whose expression is increased only in metastatic patients are underlined. The unsupervised hierarchy of each sample is shown in the top part of the figure; the unsupervised hierarchy of genes is shown on the left; the list of genes is detailed on the right. Blue=low expression, Red=high expression. **d**, Scoring of the PI3Kα activation transcriptomic signature was used to cluster patients with low, medium (both groups were combined, high+medium: red) and low (black) scoring levels in the primary tumours of confirmed PDAC patients from PAAD (TCGA) or PACA-AU (BH corrected p values). The survival curves of each cluster were then plotted, and the statistical significance was calculated using the logrank test. **e**, PI3Kα activation scores from localised PDAC in the TCGA database according to their UICC staging were retrieved, clustered into two groups (low and med) (BH corrected p values) and their overall survival was plotted, with logrank testing.

### Full annihilation of PI3Kα prevents pancreatic cancer cell migration

We then investigated on which PI3K isoform activity pancreatic cancer cells could be specifically dependent. We used a panel of murine pancreatic tumour cell lines generated by KRAS mutation combined with other genetic alterations including mutation of *PIK3CA* and deletion of *PTEN* and one commonly used human pancreatic cancer cell line PANC-1 (suppl. Table 4). We compared two different pharmacological strategies of PI3K inhibition using compounds with either isoform-selective or pan-PI3K pharmacological profiles as shown by inhibitory concentration 50 (IC50) on recombinant protein in vitro (Fig. 2a). In the 4 cell lines that we tested by WB, detectable effects on pS473Akt levels (used as read-out of PI3K activity) for all inhibitors were observed at 1μM and 10μM, thus allowing comparison of the differential downstream actions of PI3K isoforms. The most potent inhibitors, BKM120 and GDC0941 almost completely annihilated the pS473Akt levels at 10μM in the four cell lines and at 1μM in only one cell line (R211, data shown in main figure) (Fig. 2b-d, suppl. Fig. 3) suggesting that effective concentrations (EC50) are higher in pancreatic cancer cells compared to in vitro IC50. α-selective inhibitors and pan-PI3K inhibitors with low in vitro IC50 on PI3Kα significantly decreased pS473Akt levels in the four cell lines at 1 μM, suggesting that inhibitor selectivity profile is conserved.

**Figure 2:**
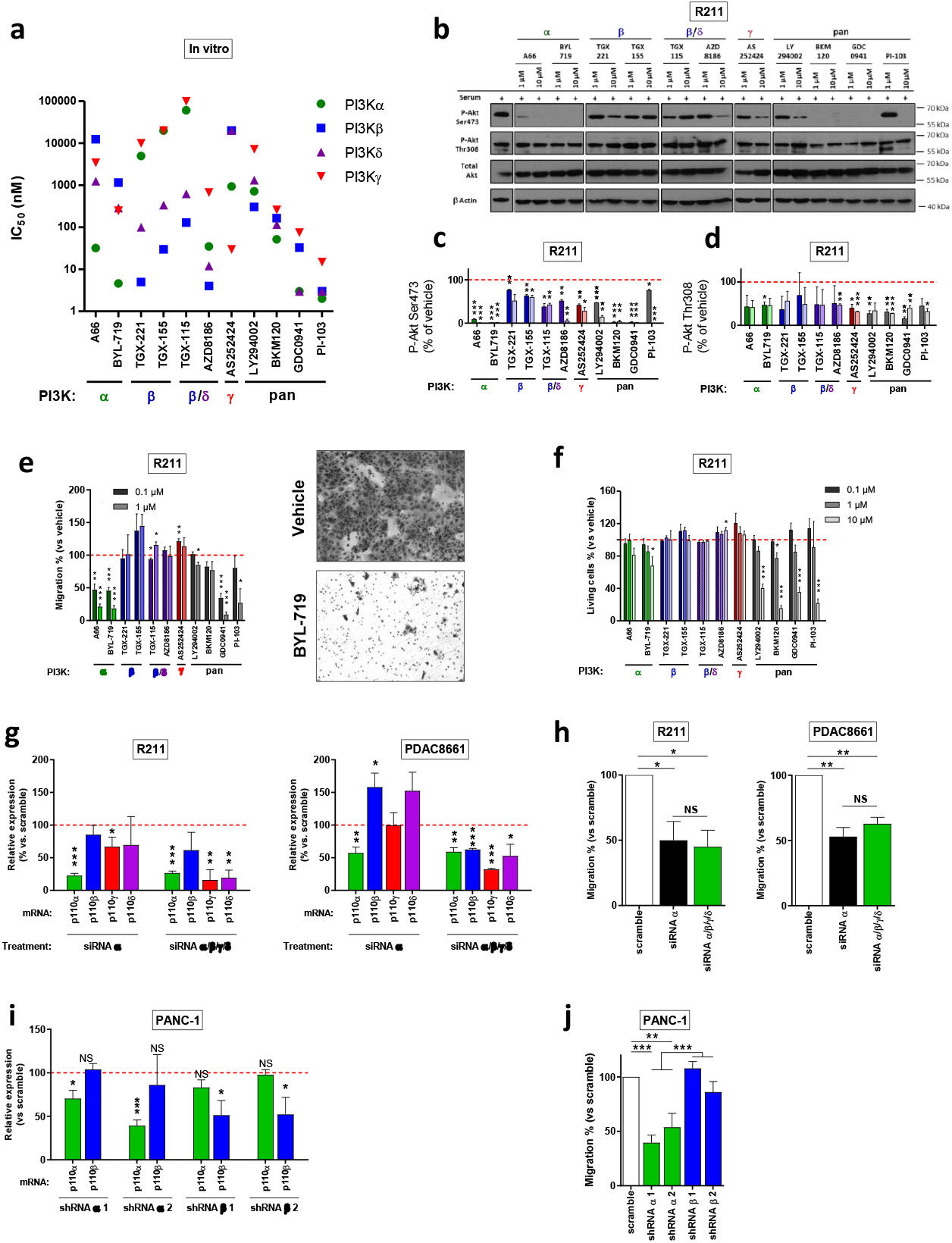
Full annihilation of PI3Kα activity is required to prevent pancreatic cancer cell migration. **a**, In vitro IC50 of PI3K inhibitors, obtained on recombinant proteins and based on the literature. The Greek letter on the x axis indicates the most potently targeted isoform. **b**, Murine pancreatic tumour cells R211 were treated for 15min with the vehicle, α, β, β/δ, γ-specific or pan-PI3K inhibitors at 1 or 10 μM in the presence of 10% FBS and the protein levels of pAkt (Ser473 and Thr308), total Akt and β Actin were observed by western blot. P-Akt on **c**, Ser473 and **d**, Thr308 were quantified and normalised with β Actin. **e**, Murine pancreatic tumour cells R211 were treated with the vehicle, α, β, β/δ, γ-specific or pan-PI3K inhibitors at 0.1 or 1 μM and simultaneously subjected to a Boyden chamber migration assay. Migrating cells were quantified after 24h. **f**, Murine pancreatic tumour cells R211 were treated with the vehicle, α, β, β/δ, γ-specific or pan-PI3K inhibitors and living cells were quantified after 3 days with a MTT colorimetric assay. Metabolically active cells are considered as living cells. **g**, Relative p110α, β, γ or δ mRNA expression (compared to scramble siRNA) after inhibition of expression by siRNA targeting p110α or a combination of siRNA targeting each class I PI3K isoform. **h**, R211 and PDAC8661 cells were treated with siRNA scramble, siRNA targeting p110α or a combination of siRNA targeting each class I PI3K isoform (pools) and subjected to a Boyden chamber migration assay. Migrating cells were quantified after 24h. **i**, Relative p110α, β, γ, or δ mRNA expression (compared to scramble shRNA stably transduced-cells) in human pancreatic tumour cells PANC-1 stably transduced with shRNA targeting p110α or β. **j**, PANC-1-transduced cells were subjected to a Boyden chamber migration assay. Migrating cells were quantified after 24h. Mean +/− SEM (* p<0.05, ** p<0.01, *** p<0.001, n ≥ 3, Student’s t-test. When not precised, comparisons are performed with vehicle).

When α-selective inhibitors, A66 and BYL-719, were used at low concentrations of 0.1 μM, significant effects were observed for assays related to migratory phenotype as shown in suppl Fig. 4,5 and in Fig. 2e-f. In all human or murine cell lines tested, A66 and BYL-719 presented a concentrationdependent capacity to inhibit pancreatic cancer cell migratory hallmarks, cell motility (suppl. Fig. 4) and directed cell migration (Fig. 2e, suppl. Fig. 4). The EC30 for migration was reached in 7/9 and 9/9 cell lines for A66 and BYL-719, respectively (Suppl. Fig. 6b). On the contrary, these parameters were not attained with the γ-specific inhibitor AS252424 and the β and β/δ-specific inhibitors except for AZD8186 which reached the EC30 for migration in 2 cell lines (suppl. Fig. 6b). The R6065 cell line (harboring *PTEN* deletion) is an exception, with two β-selective inhibitors being active on migration assays at 0.1 μM (suppl. Fig. 4b). This shows that motility and migration are pancreatic cancer cell activities sensitive to PI3Kα inhibition. Effects on cell survival/proliferation were found significant most commonly at higher concentrations (10 μM) (Fig. 2f, suppl. Fig. 5). The growth inhibitory GI30 for cytotoxicity was reached in a lower number of cell lines for both A66 and BYL-719 (Fig. 2f, suppl. Fig. 5-6a). Cell migration and cell survival of the non-tumoural ductal pancreatic cell line, HPNE hTERT, were also sensitive to PI3Kα-selective inhibition but to a lesser extent for the migration assay (suppl. Fig. 7).

To validate the pharmacological approach, we used a genetic strategy and treated R211 and PDAC8661 murine pancreatic tumour cells with pools of scramble siRNA, siRNA targeting p110α or mixed siRNA targeting each class I PI3K catalytic sub-unit, with the latter condition mimicking a pan-PI3K inhibitor. Pools of p110α-targeting siRNA or pools of p110α/β/γ/δ siRNAs lead to decreased expression of PI3K catalytic subunits (Fig. 2g) and induced similar migration inhibition on R211 and PDAC8661 cells compared to pools of scramble siRNAs (Fig. 2h). We noticed that, in PDAC8661, decreased expression of p110α mRNA increased p110β and p110δ mRNA expression, without leading to a compensatory increased migration. We also created pools of PANC-1 cells stably transfected with two hairpins targeting *PIK3CA* or two hairpins targeting *PIK3CB* (genes encoding for PI3Kα and PI3Kβ, respectively), as well as one scramble hairpin (Fig. 2i). Only shRNA targeting PI3Kα significantly decreased cell migration (Fig. 2j). Genetic approaches confirm the selective action of the PI3Kα on the migratory phenotype of pancreatic cancer cells.

### PI3Kα inhibition regulates selective PI-3,4,5-P_3_ species

To explain the downstream differences observed with PI3Kα and pan-PI3K inhibitors, we researched a concentration where both inhibitors induced the same effect on Akt phosphorylation (Fig. 3a,b). Interestingly, at this concentration, the α-selective compound presented a moderate action on the number of living cells while drastically affecting cell migration; the pan-PI3K inhibitor compound, BKM120, while being as potent on the inhibition of Akt phosphorylation, did not reduce these parameters significantly (Fig. 3c,d). This compelling result could be explained by the differential selective decrease of PI-3,4,5-P_3_ or of PIP_2_ (comprising both the substrate PI-4,5-P_2_ and the product PI-3,4-P_2_) total levels by each inhibitor, respectively (Fig. 3e). A66 as opposed to BKM120, selectively reduced the proportion of C36:2 PIP_3_ (Fig. 3f). BKM120 led to the modification of other PIP_3_ species percentages (Fig. 3f). The distributions of PIP and PIP_2_ sub-species were not modified (suppl. Fig. 8). Altogether, these data suggest that, in PDAC, PI3Kα-specific actions include production of PIP_3_ with distinctive acylation pattern, that could explain the selective promotion of cell motility and migration by PI3Kα.

**Figure 3:**
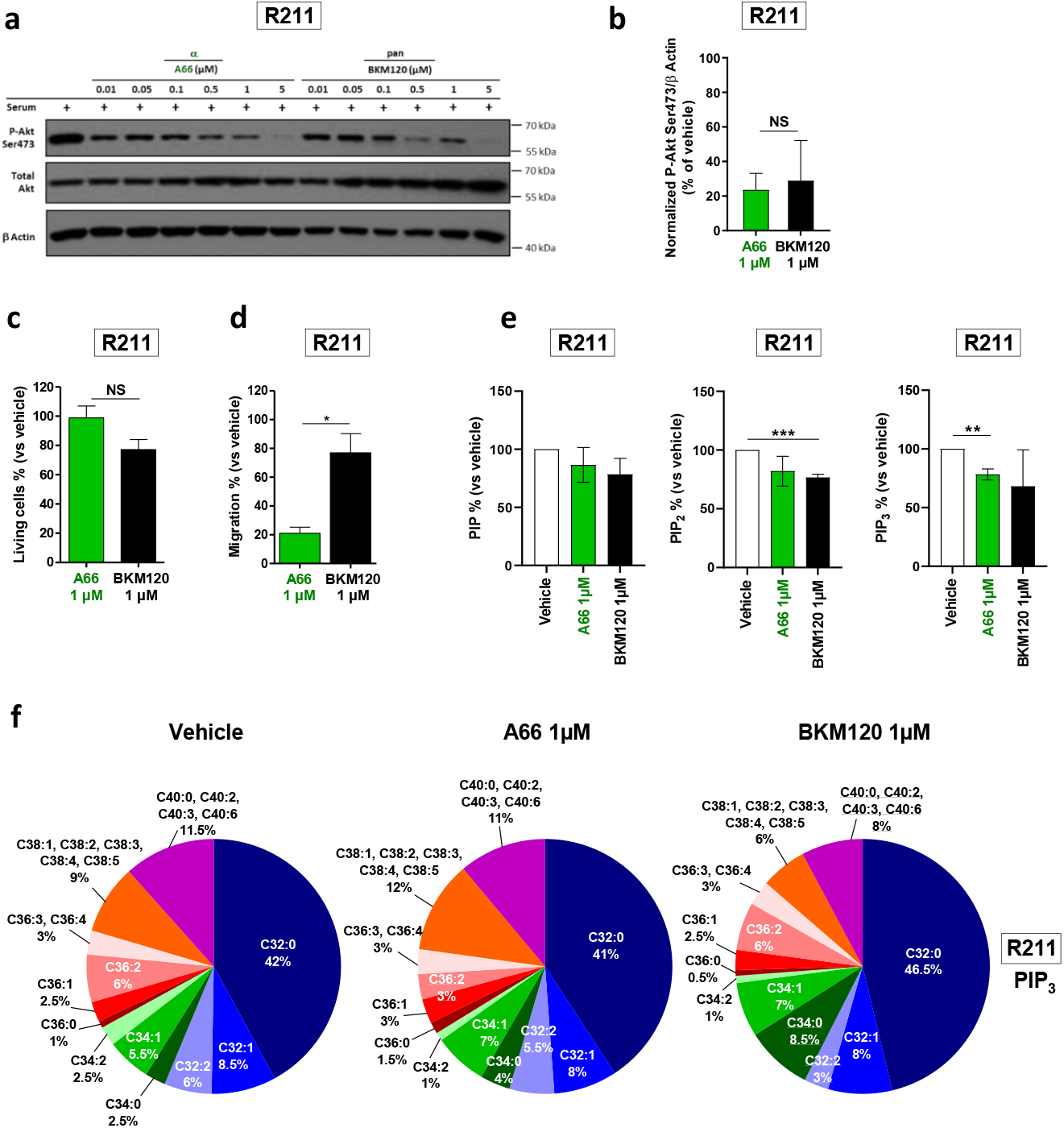
PI3Kα inhibition regulates selective PI-3,4,5-P3 species. **a**, Murine pancreatic tumour cells R211 were treated for 15 min with the vehicle, α-specific A66 or pan-PI3K inhibitor BKM120 at 0.01, 0.05, 0.1, 0.5, 1 or 5 μM in the presence of 10% FBS and protein levels of P-Akt (Ser473), total Akt and β Actin were observed by western blot. **b**, P-Akt on Ser473 was quantified and normalised with β Actin. R211 cells were treated with the vehicle, α-specific inhibitor A66 or pan-PI3K inhibitor BKM120 at 1 μM and **c**, living cells were quantified after 3 days with a MTT colorimetric assay or **d**, migrating cells were quantified after 24h in a Boyden chamber assay and compared to vehicle (DMSO). **e**, Murine pancreatic tumour cells R211 were treated for 15 min with the vehicle (0.01% DMSO) or α-specific A66 or pan-PI3K inhibitor BKM120 at 1μM. Phospholipids were extracted, total PIP, PIP_2_ and PIP_3_ were quantified and compared to the vehicle (DMSO). **f**, the proportion of each PIP3 sub-type was represented. Sub-types whose proportion was modified compared to DMSO were underlined. Mean +/− SEM (* p<0.05, ** p<0.01, *** p<0.001, n ≥ 3, Student’s t-test).

### PI3Kα inhibition targets cell migration and cell survival regardless of the genetic landscape of pancreatic adenocarcinoma cells

PI3Kα oncogenic action is commonly described as directly coupled only to oncogenic KRAS or tyrosine kinase receptors, and not to *PTEN* alterations (20). To challenge this concept, we analysed the impact of the genetic alterations (on *KRAS, PIK3CA, PTEN*) and of the organ of origin in determining the role of each class I PI3K in tumour cell migration and cytotoxic sensitivity in response to PI3K inhibitors. With the data shown in Fig. 2, suppl. Fig. 4e-f, suppl. Fig. 5d-e, we calculated the correlation between the in vitro IC50 of PI3K inhibitors for all class I PI3Ks (Fig. 2a, suppl. M&M), and their capacity to inhibit migration in 9 cell lines (correlation test values are shown in Fig. 4e-f, individual values for each cell lines in suppl. Fig. 9). The ability of all PI3K inhibitors (at 1μM) to regulate cell migration mainly depended on their capacity to target PI3Kα, and in a less frequent way on PI3Kβ, δ or γ (Fig. 4a, suppl. Fig. 9). We reported the p-value of the effect vs. IC50 correlation test for each class I isoform and found that PI3Kα was significantly associated with pancreatic cell migration (Fig. 4b, suppl. Fig. 9c), cell motility (suppl. Fig. 9a,e), cytotoxicity (Fig. 4c, suppl. Fig. 9d) and pSer473Akt phosphorylation (suppl. Fig. 9b,f). As a positive control, we confirmed the demonstrated isoform dependency that was published by others in other solid and liquid cancer cell lines (21–23). Most cancer cell lines of pancreatic origin depended on PI3Kα activity, and sometimes on PI3Kδ and PI3Kγ to regulate their migration and cell viability despite the genetic context. This result suggests that PI3Kα would be a target of choice for pancreatic cancer patients who, despite their genetic heterogeneity associated with class I PI3K activation (mutant *KRAS*, deletion of *PTEN*, mutation of *PIK3CA*), are all dependent on intrinsic basal PI3Kα activity, thus corroborating the prognostic value of the PI3Kα activity signature to detect pro-metastatic features (Fig. 1).

**Figure 4:**
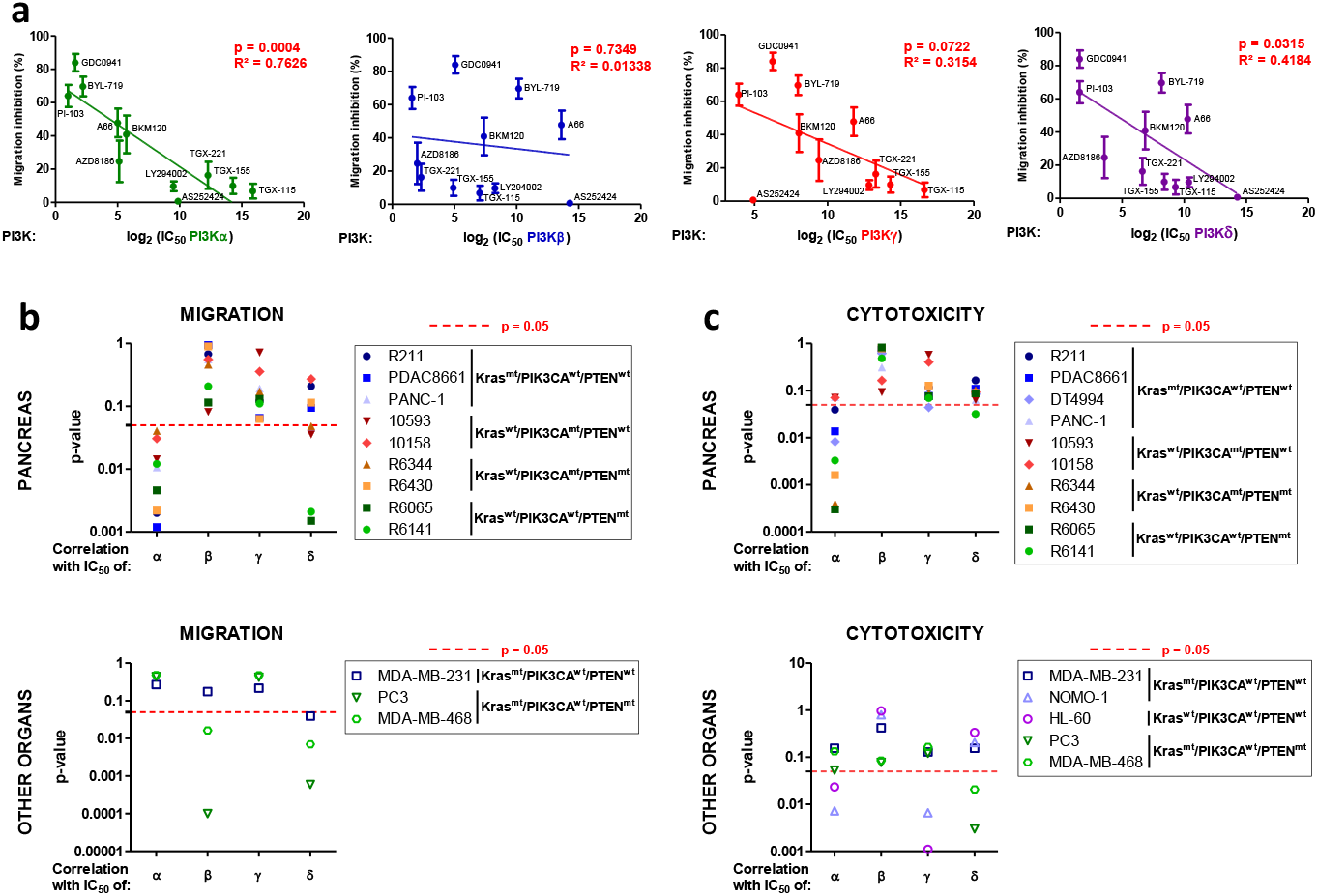
PI3Kα inhibition is effective on pro-tumoural features regardless of the genetic landscape of pancreatic adenocarcinoma. **a**, Migration inhibition capacities of PI3K inhibitors at 1 μM were tested on pancreatic tumour cells (R211, PDAC8661, PANC-1, 10593, 10158, R6344, R6430, R6065, R6141). The mean of these values was plotted against the in vitro IC50 for each class I PI3K and each inhibitor. IC50 is determined on recombinant proteins. Pearson correlation tests were performed and p-values were presented separately for each class I PI3K isoform. **b**, individual p-values of these correlation tests were represented for each of the 9 pancreatic (R211, PDAC8661, PANC-1, 10593, 10158, R6344, R6430, R6065, R6141,) breast (MDA-MB-231, MDA-MB-468) and prostate (PC3) cancer cell lines; values for each PI3K isoforms are plotted separately. **c**, Cytotoxic capacities of PI3K inhibitors at 10 μM were tested on the 9 pancreatic (R211, PDAC8661, PANC-1, 10593, 10158, R6344, R6430, R6065, R6141), breast (MDA-MB-231, MDA-MB-468), prostate (PC3) and acute myeloid cells (NOMO-1, HL-60) and the correlation with the in vitro IC50 for each class I PI3K of each inhibitor was determined. Pearson correlation tests were performed and p-values represented for each isoform and cell line. The dotted red line corresponds to a threshold p value of 0.05. Mean +/− SEM (* p<0.05, ** p<0.01, *** p<0.001, n≥3, Student’s t-test).

### Pharmacological PI3Kα inhibition prevents the rapid progression of cfDNA-positive PDAC

While PDAC patients with increased levels of ctDNA present a worse prognosis (16, 17), cfDNA and ctDNA could be indicative of underlying micrometastatic disease. When we compared patients with and without pathological nodal involvement (pT1N0 versus pT2N1), we observed a stronger pAkt-Substrate IHC staining indicative of PI3K/Akt activity in the primary tumour/pancreas and a higher level of cfDNA with detected KRAS mutation on pT2N1 stage, which suggest a correlation with dependency to PI3K activity and metastatic potential (Figure 5a).

**Figure 5:**
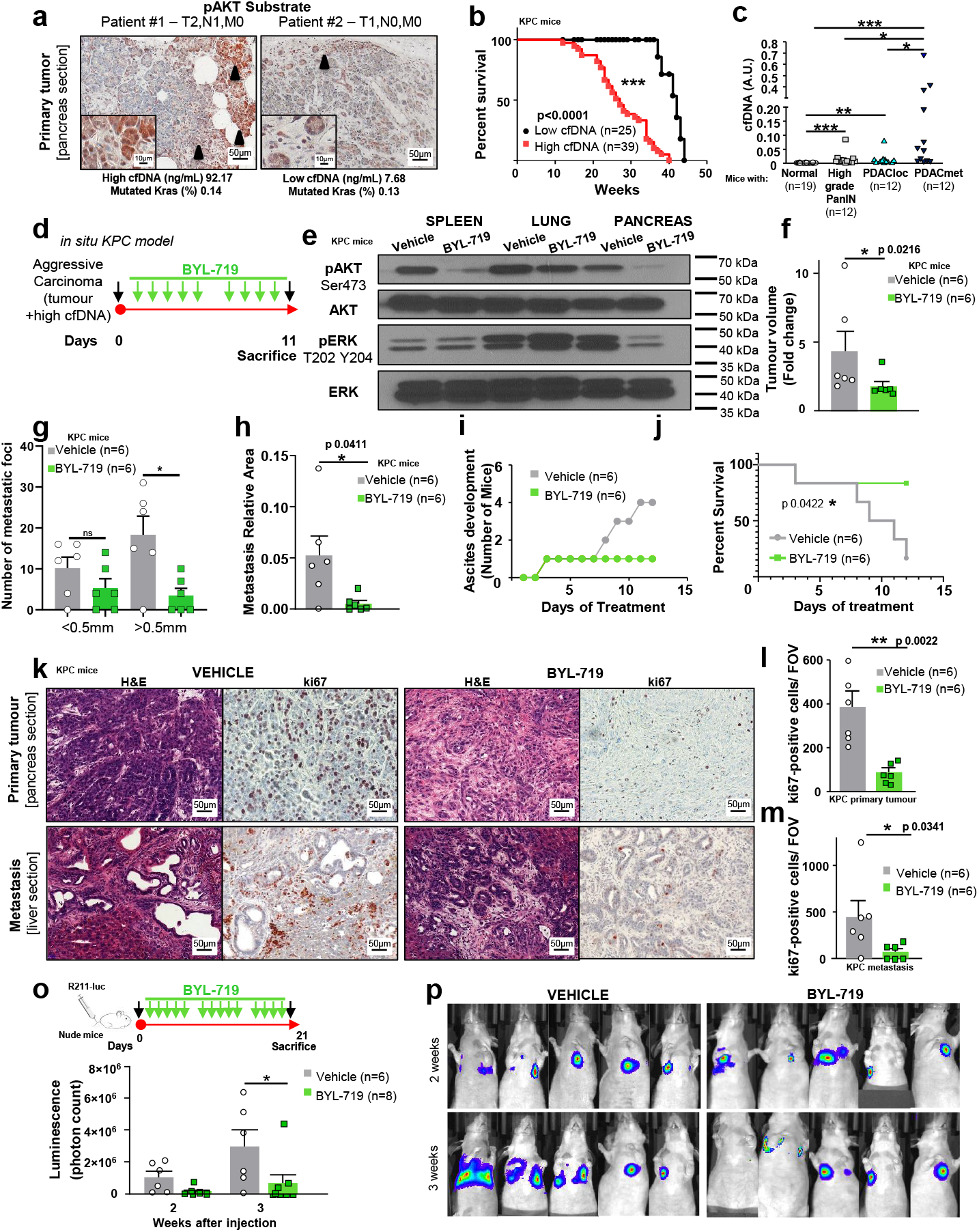
Pharmacological PI3Kα inhibition prevents the rapid progression of cfDNA-positive PDAC. **a,** Representative images of PI3K activation (as shown by positive pAkt substrate staining) on the primary tumours of these PDAC patients, with values of quantification of cfDNA and KRAS allele frequency. **b**, Survival curve of KPC mice presenting different levels of cfDNA. N in each group is indicated. **c,** Quantification of cfDNA in KPC mice at different stages of the disease. N in each group is indicated. **d**, KPC mice diagnosed with an aggressive carcinoma were given daily oral doses of BYL-719 (50mg/kg). n=6 in each group **e**, The protein levels of P-Akt (Ser473), total Akt, pERK (Thr202 and Tyr204) and total ERK were observed by western blot in spleen, lung and pancreas tissue lysates. **f**, Quantification of tumour volume (expressed in fold change) and **g-h**, quantification of micro- and macro-metastatic foci and the relative metastasis area (comprising metastases in the spleen, liver and lung) of KPC mice treated with the vehicle and BYL-719. **i**, Development of ascites in KPC mice. **j**, Survival curve of KPC mice treated with the vehicle or BYL-719. **k**, Representative images and **l,** quantification of Ki67 positive cells in pancreatic tumours and in **m**, metastasized liver sections. **o**, Treatment regimen of the tail vein injection experiment in nude mice treated with the PI3Kα inhibitor or vehicle (n=6 in each group), quantification at two time points and **p**, representative images of luminescence via IVIS^®^ Spectrum in vivo imaging system. Mean +/− SEM (* p<0.05, ** p<0.005, *** p<0.0001)

We measured cfDNA in the KPC model to test the effects of PI3Kα-selective pharmacological inhibition on established tumours presenting these circulating markers of metastatic disease. In the KPC model, aggressive pancreatic tumours spontaneously develop under KRAS and p53 oncogenic mutations (24), however detection of cfDNA has not been tested yet. Aggressive tumours were diagnosed through high-resolution ultrasound (US) imaging as well as quantification of cfDNA (for the setting of threshold limit, see also suppl. M&M). We quantified the cfDNA in blood plasma samples by longitudinally measuring the relative concentration of two expressed genes, *TP53* and *GAPDH* genes and then correlated those findings with the anatomo-pathological results from the pancreas and metastatic site organs (suppl. Table 5). The longitudinal average levels of cfDNA correlated with disease progression and mouse lethality (Fig. 5b). We were also able to predict the pathology (localised primary tumour vs. metastasis detection) by the level of cfDNA at sacrifice in two independent cohortes (suppl. Fig. 10a-e). Short fragment of cfDNA was shown by others to be specific to tumour cells (25). Analysis of cfDNA integrity revealed a distinct 160-210bp fragment selectively increased in mice that developed metastatic PDAC compared to mice with high-grade PanINs and localised PDAC (suppl Fig. 10b).

We then treated KPC mice featuring aggressive carcinoma (i. e. featuring a echography-detected tumour and a high level of cfDNA, suppl. Fig. 11), with the PI3Kα-selective inhibitor, BYL-719, or with the vehicle (Fig. 5d) (26). pS473-Akt levels were reduced by BYL-719 treatment in all tested tissues, while pERK-T202 Y204 levels fell only in the pancreas following BYL-719 treatment (Fig. 5e). We stopped the cohort treatment when all the vehicle treated mice presented markers of macroscopic metastatic dissemination determined by ascites detection or ethical limit points. This was done in an aim to explain the mechanisms responsible for BYL-719 action. The BYL-719 treatment line significantly slowed tumour volume progression (Fig. 5f, suppl. Fig. 12) and reduced the number of macrometastatic foci (Fig. 5g), distant metastatic area in lung, liver and spleen (Fig. 5h), delayed ascites development as a marker of dissemination (Fig. 5i) and mice survival (Fig. 5j). While all the vehicle mice had to be sacrificed during the treatment only one BYL-719-treated mouse presented signs of tumour evolution that necessitated immediate sacrifice. A correlation was established between the proliferative index of cancer cells (assessed by ki67 index in primary tumour and metastatic sites), which reflects the global tumour burden, and levels of cfDNA (suppl. Fig. 13a). BYL-719 significantly reduced cell proliferation assessed by Ki67 index in both primary and metastatic sites (Fig. 5k-m, suppl. Fig. 13b, with associated Power test in suppl Table 5), associated with decreased tumour grading and changes in tumour cell/stroma ratios (Suppl. Fig. 14). The anticipated secondary effects of BYL-719 treatment on insulin secretion were observed (suppl. Fig. 15a,b). We did not see any significant impact on levels of cleaved-caspase 3 positive cells (apoptotic marker: suppl. Fig. 15c) or on γ-H2AX positive foci (DNA damage marker: suppl. Fig. 15d).

Furthermore, in a context of early pancreatic cancer lesions induced by oncogenic KRAS in an inflammatory condition (caerulein injections), the highly selective pharmacological inactivation of PI3Kα with another compound, namely GDC-0326, completely prevented the maintenance of precancer lesions (suppl. Fig. 16 a-d), and features linked to stromal remodelling (Suppl. Fig. 16b,e). The high level of cleaved caspase 3 detected in epithelial lesions could be prevented by PI3Kα inactivation (suppl. Fig. 16b,f).

Tail vein injection of murine pancreatic cancer cells, engineered to express secreted luciferase (R211-Luc cells) in Nude mice, was carried out in order to confirm the action of BYL-719 treatment on evolution of micrometastatic foci. Administration of BYL-719 treatment for 21 days significantly prevented tumour cell growth in lung (Fig 5. l,m, suppl. Table 5). Taken together, these data demonstrate that in vivo inhibition of PI3Kα prevents the evolution of micrometastatic foci into macrometastatic foci and the rapid progression of aggressive cfDNA-positive PDAC.

### Tumour-intrinsic PI3Kα alters tumour cell chemokine secretion and promotes the acquisition of protumoural M2 macrophage characteristics in the peritumoural tissue

We performed a full blood count to assess immune cell populations (suppl M&M, Suppl. Table 5), as macrophages promote PDAC progression (27). We noticed that metastatic KPC mice presented significantly increased counts of white blood cells (Fig. 6a), monocyte (Fig. 6b) and granulocyte (Fig. 6c) counts compared to KPC with localised tumours. The lymphocyte counts remained the same (Fig. 6d).

**Figure 6:**
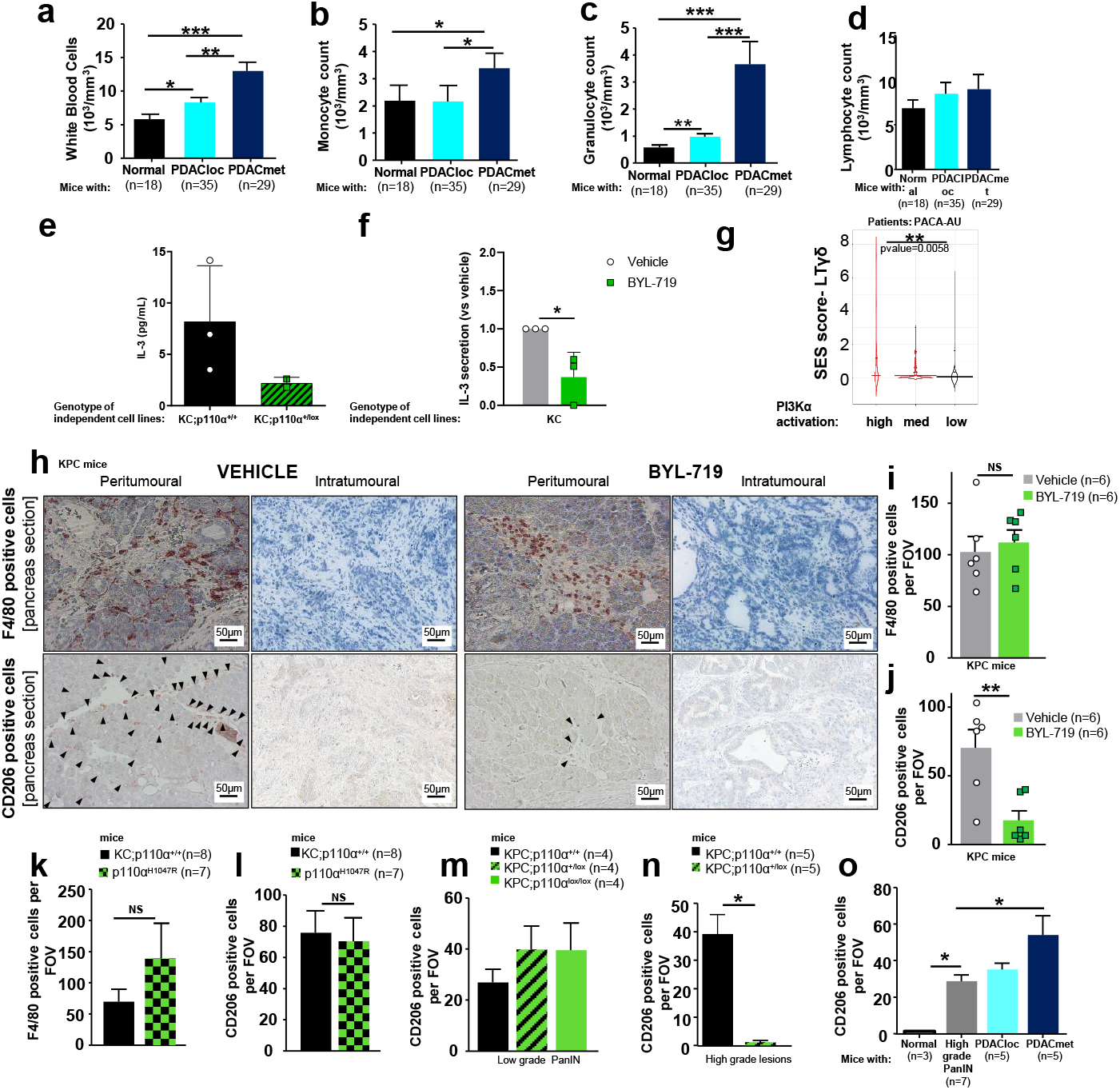
Tumour-intrinsic PI3Kα alters tumour cell chemokine secretion and promotes the acquisition of protumoural M2 macrophage characteristics in the peritumoural tissue. Blood count of **a,** white blood cells, **b,** monocytes, **c,** granulocytes and **d,** lymphocytes in KPC mice, with cohort size in each group. **e,** Basal level of IL-3 was quantified in KC;p110α^+/+^ (R211, A338 and A338L) and KC;p110α^+/lox^ (A260 and A94L) cell lines isolated from mice primary tumours or lung metastases (L). **f,** IL-3 relative level (compared to vehicle) was determined in 3 KC (R211, A338 and A338L) cell lines isolated from mice primary tumours under treatment with vehicle (DMSO) or 1μM of BYL-719. **g,** Violin plot demonstrating the link between PI3Kα activation and activation of a selective population of γδLT in the PAAD cohort. **h,** Representative images and quantification of **i,** F4/80 positive cells and **j,** CD206 positive cells after the pharmacological inhibition of PI3Kα. Quantification of **k,** F4/80 and **l**, CD206 positive cells in KC and oncogenic PI3Kα mice. Quantification of CD206 positive cells in **m,** low-grade and **n,** high-grade PanIN lesions after the genetic inactivation of PI3Kα in epithelial pancreatic cells. **o,** Quantification of CD206 positive in KPC mice at different stages of PDAC progression. N in each group is indicated in the figure. Mean +/− SEM (* p<0.05, ** p<0.005, *** p<0.0001).

To test whether tumour-intrinsic PI3Kα could be involved in this tumour/stroma interaction, we derived cell lines from tumoural lesions induced either by Kras mutant or Kras mutant in pancreatic cells partly lacking PI3Kα activity (see also M&M). We researched and found that genetic inactivation of PI3Kα in pancreatic cancer cells led to an altered cytokine secretion pattern in vitro (a panel of 16 cytokines were tested), with levels of IL-3 decreased in 2 mutant KRAS pancreatic cancer cell lines partly lacking PI3Kα activity (genetic inactivation) compared to 3 mutant KRAS pancreatic cancer cell lines including R211 cells (Fig. 6E, suppl. Fig. 17). Pharmacological inactivation of PI3Kα in 3 different mutant KRAS pancreatic cancer cell lines decreased significantly and reproducibly IL3 levels (Fig. 6f). PDAC patients with a high PI3Kα activity signature presented a significant increase of the gene signature of the selective immune population of γδ T lymphocytes (LTγδ) and not of the other immune cell population generic signatures (Fig. 6g, suppl. Table 6). This immune cell signature is known to be associated with poor prognosis and highly inflammatory conditions, promoting differential macrophage differentiation (28).

We therefore analysed immune cell composition in the cohort of KPC mice treated with BYL-719 or the vehicle. BYL-719 did not modify the overall F4/80^+^ macrophage counts (Fig 6h,i), but significantly prevented their differentiation into pro-tumourigenic CD206^+^ macrophages (M2 macrophages) in tumour adjacent tissues (Fig. 6h,j), with no significant difference in CD4 and CD8 infiltration (suppl. Fig. 18). To test whether oncogenic PI3Kα mimics oncogenic Kras also on tumour/stroma interaction, F4/80^+^ macrophage recruitment and infiltrating CD206-macrophages around tumours were quantified and found at a similar rate in peritumoural tissue from hyperactivated oncogenic PI3Kα (p110α^H1047R^) and KRAS-mutant tumours (Fig. 6k,l); oncogenic PI3Kα did not trigger inflammation in normal pancreas of young mice (1-3 months old) as assessed by quantifying infiltrating F4/80 macrophages (suppl. Fig. 18b). Inactivation of PI3Kα only in pancreatic epithelial cells was sufficient to completely prevent M2 macrophage infiltration selectively around high grade lesions (Fig. 6m,n, suppl. Fig. 18c,d), in favour of tumour cell intrinsic action of PI3Kα. Finally, increased peritumoural CD206 staining is associated with further development of metastatic foci and ascites (Fig. 6o, suppl. Fig. 18e).

Our data show that the rapid progression of aggressive cfDNA-positive PDAC driven by PI3Kα involves changes in the inflammatory cytokine context in conjunction with M2 pro-tumoural macrophage characteristics in peritumoural tissue.

## Discussion

In the path of the recent success of Olaparib as a targeted therapy in advanced pancreatic metastatic patients (29, 30), further attempts should be made to prevent their rapid progression. We have demonstrated that one such strategy could be to target the PI3Kα-driven signal that critically sustains several aspects of pancreatic oncogenicity, including those at the origin of tumour-induced environment rewiring and metastasis evolution (**Graphical Abstract**).

In the clinical setting, single agents targeting PI3K had a limited impact (31), which was most probably due to the non-optimised selection of patients with advanced disease (32). Interestingly, in metastatic breast cancer (MBC), when *PIK3CA* mutation was detected in circulating tumour DNA, patient disease-free survival (PFS) improvement became largely significant (33), and BYL-719 (alpelisib) was recently granted marketing authorisation for the treatment of breast cancer in combination with endocrine therapy (26). Our study suggests that contrary to MBC, even without *PIK3CA* oncogenic mutation, PI3K pathway could be a driver of the pancreatic metastatic evolution, and thus a druggable target. In this cancer setting, high levels of circulating tumour DNA as a marker of micrometastatic disease were found to be correlated with poor prognosis (16). Experimental data from other investigators also argue that disseminating cells are detected very early in the disease history (34). Our longitudinal analysis of circulating DNA levels demonstrates an early increase of these parameters in an experimental pancreatic cancer model. Measurement of the concentration of the 160-210 bp fragment increases the prediction rate of survival, as evidenced in patients in other studies using circulating mutation rates (35). In terms of toxicity, the main concern for using potent PI3Kα inhibitors remains the induction of insulin feedback which could feed the tumours (2). These concerns could be resolved by the clinical management of glycaemia during treatment (2, 36). It has to be noted that this insulin feedback occurs to be reduced by age and that PDAC is mostly detected in patients >40 years (37).

Our data also demonstrate that oncogenic KRAS-PI3Kα coupling leads to specific functions of clinical relevance in the pancreatic context. In mutant KRAS-driven lung cancers, inactivation of PI3Kα yielded only a partial response (38), MAPK pathway activity being key in this particular context. From our results, there are four non-exclusive explanations of the importance of KRAS-PI3Kα coupling in this organ setting.

Firstly, in the pancreas, PI3Kα selective inactivation induced decreased phosphorylation of Erk1/2. Our data are not in line with data from other studies in this regard (39–41); other authors used pan-PI3K inhibitors. MEK inhibition has a minimal anti-tumoural action due to the induction of strong feedback activation on Akt (42). Selectivity towards PI3Kα could prevent the induction of compensatory signals towards the MAPK pathway.

Secondly, the action of PI3Kα on actin cytoskeleton remodelling in pancreatic cancer cells is very sensitive to pharmacological inhibition, regardless of genetic background. We can speculate that remodelling of actin cytoskeleton appears to be key in PDAC progression so that tumour cells can extrude themselves from the strong desmoplastic and low vascularised pancreatic primary tumours. Very early on in the discovery of oncogenic KRAS properties, it was demonstrated that the PI3K signal towards actin remodelling was the first major signalling event leading to cell transformation (43).

Subsequently, it was also found that actin remodelling was linked to PI3Kα-driven glucose metabolism regulation (44). Actin cytoskeleton remodelling and associated cellular functions such as migration and motility are sensitive to PI3Kα inhibition. Currently, published data are increasingly showing that the acylation state of phospholipids could modulate their localisation and function (45). In this context, we can provide additional information to show that pan-PI3K inhibitors or isoform-specific inhibitors could target different PIP_3_ subspecies and conversion to PI-3,4-P_2_ ultimately leading to clear-cut effects on migration or cytotoxicity.

Thirdly, protumoural macrophages in the organ around the tumour could favour metastasis evolution. Others found that activation of macrophages associated with increased apoptotic tumour cells accelerated growth and not implantation of prostate metastatic nodules (46). Similarly, systemic secretions of the M2 macrophages that are found around the primary pancreatic tumours could favour growth of metastatic micro-metastatic foci. Tumour-intrinsic PI3Kα indirectly rewires immune cell composition in the tumour site; this increase of CD206-positive cell counts in the tissue could act as a distant event and promote metastatic evolution. Indeed, suppression of macrophages in the KPC mice and their primary tumours by clodronate treatment did not delay lethality, but decreased incidence of macro-metastasis (47). The tumour-intrinsic action of PI3Kα on tumour inflammation that we described here could be added to its direct immunomodulatory action in pancreatic cancer found by others (48).

Fourthly, the repartition of each KRAS mutation is different in each solid cancer, with G12D mutation being the most common in PDAC. Recent evidence shows that each KRAS mutation drives different signalling and engage different pathways (49, 50). This differential engagement could explain the prominent role of PI3Kα downstream KRASG12D in PDAC. However, the multiple PI3K isoform engagement could also explain why some tumours appear to escape from BYL-719 treatment in vivo. Interestingly, while all BYL-719 treated tumours presented a significant decrease in CD206 staining, the effects on CD4 and CD8 immune cell population were found heterogeneous; this could explain why some BYL-719 treated mice presented (albeit in a reduced size and number) macro-metastatic foci. In the Nude mice model, that is devoid of lymphocytes but presents macrophages, 3 mice out of 8 also displayed a lower effect of BYL-719 treatment, suggesting that other parameters than immune cell rewiring could be involved in resistance to treatment. In vitro, some cell lines were also sensitive to other PI3K isoform inhibitor. Both PI3Kγ and PI3Kα are important for pancreatic cancer (51); the possible crosstalk between these two isoforms should be investigated. We tested in vitro whether pharmacological inhibition of PI3Kα could be synergistic with gemcitabine, the current treatment of PDAC patients, and found this was not the case (data not shown). In a future study, we aim to dissect what could be the mechanisms of resistance to PI3Kα inhibition, in an aim to increase the efficiency of this therapeutic agent on PDAC evolution.

In conclusion, our data demonstrate that PI3K-targeting agents could be effective in the management of micro-metastatic disease (measured by cfDNA), notably for PDAC patients, resulting in a delay in macro-metastatic evolution via immunomodulatory action in the peritumoural tissue.

## Methods

### Samples and animal models

Human PDAC samples and cells were collected according to French and European legislation, and stored accordingly in CRB, Toulouse, IUCT-0. All animal procedures were conducted in compliance with the Ethics Committee pursuant to European legislation translated into French Law as Décret 2013-118 dated 1st of February 2013 (APAFIS 3601-2015121622062840). General methods were used and are detailed in the supplementary material and methods.

## Acknowledgments

We are grateful to SigDYN members, past and present, for their technical support, sample banks, common tools, scientific and protocol discussions, CRB and BACAP consortium for patient samples, Anexplo team work (mouse breeding & experimental zone; ENI core platform), the CRCT core technology platform in particular Laetitia Ligat, Carine Valle and Emeline Sarot, ImagIN platform (FX Fresnois), Dr. Barbara Garmy-Susini for access to her laboratory whilst moving our own facilities, the staff of MetaToul-Lipidomique Core Facility (I2MC, Inserm 1048, Toulouse, France), MetaboHUB-ANR-11-INBS-0010 for lipidomic analysis, advices and technical assistance, and Genentech for GDC0326. JGG is a member of COST action EU-Pancreas BM1204. JGG’s laboratory belongs to Toucan, Laboratoire d’Excellence, ANR, an integrated research program on Signal-targeted Drug Resistance. JGG’s laboratory for this topic was/is funded by Europe EU-ERG FP7 (270696 PaCa/PI3K), ARC (PJA20171206596; salary for RB), Toucan ANR Laboratory of Excellence, MSCA-ITN/ETN PhD-PI3K (Project ID: 675392, salary for FRD and SA), Fondation de France (salary for BT), GSO, Ligue Nationale Contre le Cancer (salary for CC and CC).

---

**Supplementary Figure 1: PI3Kα activation signature.** Identification of PI3Kα activation signature

**Supplementary Figure 2: Distribution of genetic alterations in PAAD cohort (most frequent alterations and alterations related to PI3K activation).**

**Supplementary Figure 3: All data related to western Blot of Akt. a,** Murine (PDAC8661, DT4994) and human (PANC-1) pancreatic tumours cells were treated for 15min with α, β, β/δ, γ-selective or pan-PI3K inhibitors at 0.1 or 1 μM in the presence of 10% FBS and the protein levels of P-Akt (Ser473 and Thr308), total Akt and β Actin were observed by western blot. P-Akt on **b,** Ser473 or **c,** Thr308 was quantified and normalised with β Actin.

**Supplementary Figure 4: All data relating to migration assays. a,** Scratching was applied to a monolayer of murine pancreatic tumour cells R211 or PDAC8661. The cells were treated concomitantly with α-selective or pan-PI3K inhibitors at 0.01, 0.1 or 1 μM and observed after 24 h to determine the number of wounds that had healed. **b,** murine pancreatic cells PDAC8661, whether or not treated with BYL-719 at 1 μM were counted 24 h after wound healing assay. **c,** Murine (PDAC8661) and human (PANC-1) pancreatic tumour cells were treated with α, β, β/δ, γ-selective or pan-PI3K inhibitors at 0.1 or 1 μM. **d,** KRAS^wt/^PIK3CA^mt^ murine pancreatic tumour cells 10593 and 10158 were treated with α, γ-selective or pan-PI3K inhibitors at 0.1 or 1 μM. **e,** Murine pancreatic tumour cells (R6344, R6430, R6065, R6141) were treated with α, β, β/δ, γ-selective or pan-PI3K inhibitors at 0.1 or 1 μM. **f,** Prostate (PC3) and breast cancer cells (MDA-MB-231, MDA-MB-468) were treated with α, β, β/δ, γ-selective or pan-PI3K inhibitors at 0.1 or 1 μM. **c-f**, Cells were concomitantly subjected to a Boyden chamber migration assay and migrating cells were quantified after 24h. Mean +/− SEM (* p<0.05, ** p<0.01, *** p<0.001, n≥3, Student’s t-test).

**Supplementary Figure 5: All data relating to cytotoxicity assays. a,** Murine (PDAC8661, PDAC8661, DT4994) and human (PANC-1) pancreatic tumour cells were treated with α, β, β/δ, γ-selective or pan-PI3K inhibitors for 3 days. **b**, KRAS^wt^/PIK3CA^mt^ murine pancreatic tumour cells 10593 and 10158 were treated with α, γ-selective or pan-PI3K inhibitors. **c,** Murine pancreatic tumour cells (R6344, R6430, R6065, R6141) were treated with α, β, β/δ, γ-selective or pan-PI3K inhibitors. **d,** Acute myeloid leukaemia (HL-60, NOMO-1), prostate (PC3) and breast cancer cells (MDA-MB-231, MDA-MB-468) were treated with α, β, β/δ, γ-selective or pan-PI3K inhibitors. **a-d**, Living cells were quantified after 3 days with a MTT colorimetric assay and metabolically active cells are considered as living cells. Mean +/− SEM (* p<0.05, ** p<0.01, *** p<0.001, n≥3, Student’s t-test).

**Supplementary Figure 6: GI30 and IC30 for cytotoxicity and migration experiments. a,** Pancreatic cancer (R211, PDAC8661, DT4994, PANC-1, 10593, 10158, R6344, R6430, R6065, R6141), pancreatic untransformed (HPNE hTERT), prostate cancer (PC3), breast cancer cells (MDA-MB-231, MDA-MB-468) and acute myeloid leukaemia cells (NOMO-1, HL-60) were treated with 0.1, 1 or 10 μM of α, β, β/δ, γ-specific or pan-PI3K inhibitors and living cells were quantified after 3 days with a MTT colorimetric assay. GI30 (for the concentration that inhibited 30% of cell growth) was determined with GraphPad Prism 8 using log (inhibitor) vs. response - variable slope (four parameters) analysis. “-” indicates that the inhibitor was not used in the corresponding cell line, “NR” indicates that the GI30 was not reached. **b,** Pancreatic cancer (R211, PDAC8661, DT4994, PANC-1, 10593, 10158, R6344, R6430, R6065, R6141), pancreatic untransformed (HPNE hTERT), prostate cancer (PC3) and breast cancer cells (MDA-MB-231, MDA-MB-468) were treated with 0.1 or 1 μM of α, β, β/δ, γ-specific or pan-PI3K inhibitors and concomitantly subjected to a Boyden chamber migration assay. Migrating cells were quantified after 24h. CI30 (for the concentration that inhibited 30% of cell migration) was determined with GraphPad Prism 8 using log (inhibitor) vs. response - variable slope (four parameters) analysis. indicates that the inhibitor was not used in the corresponding cell line, “NR” indicates that the GI30 was not reached. Mean +/− SEM (* p<0.05, ** p<0.01, *** p<0.001, n≥3, Student’s t-test).

**Supplementary Figure 7: HPNE hTERT cell cytotoxicity and migration experiments. a,** Ductal pancreatic cells HPNE hTERT were treated with α, β, β/δ, γ-specific or pan-PI3K inhibitors and living cells were quantified after 3 days with a MTT colorimetric assay. Metabolically active cells are considered as living cells. **b,** Cytotoxic capacities of PI3K inhibitors at 1 μM were tested on HPNE hTERT cells and were plotted against the in vitro IC50 for each class I PI3K of each inhibitor determined on recombinant proteins. Pearson correlation tests were performed. **c,** HPNE hTERT cells were treated with α, β, β/δ, γ-specific or pan-PI3K inhibitors at 0.1 or 1 μM and concomitantly subjected to a Boyden chamber migration assay. Migrating cells were quantified after 24 h. **d,** Migration inhibition capacities of PI3K inhibitors at 1 μM were tested on HPNE hTERT cells and were plotted against the in vitro IC50 for each class I PI3K of each inhibitor determined on recombinant proteins. Pearson correlation tests were performed. Mean +/− SEM (* p<0.05, ** p<0.01, *** p<0.001, n≥3, Student’s t-test).

**Supplementary Figure 8: PIP and PIP_2_ sub-types.** Murine pancreatic tumour cells R211 are treated for 15 min with the vehicle (0.01% DMSO) or α-specific A66 or pan-PI3K inhibitor BKM120 at 1μM. Phospholipids were extracted and the proportion of each **a,** PIP and **b,** PIP2 subtype was represented. N=3

**Supplementary Figure 9: Correlations between cellular phenotypes and class I PI3K isoform inhibition.** Inhibitory capacities of PI3K inhibitors at 1 μM on motility and at 10 μM on Akt phosphorylation (Ser473) were plotted against the in vitro IC50 for each class I PI3K (α, β, γ and δ) of each inhibitor determined on recombinant proteins. Pearson correlation tests were performed. The p-value obtained was represented for each tested cell line and for each class I PI3K isoform for **a,** motility and **b,** Akt phosphorylation on Ser473. The dotted red line corresponds to a p-value of 0.05. Individual R squared values of the Pearson correlation test for each cell line (from the pancreas or other organs) and each PI3K isoform are shown for **c,** migration, **d,** cytotoxicity, **e,** motility and **f,** Akt phosphorylation on Ser473. Mean +/−SEM (* p<0.05, ** p<0.01, *** p<0.001, n≥3, Student’s t-test).

**Supplementary Figure 10: Additional information on cfDNA in KPC mice. a,** Quantification of cfDNA by GAPHD gene in KPC mice at different stages of the disease. **b,** Quantification of 160-210bp Fragment concentration in KPC mice at different stages of the disease. N in each group is indicated. **c,** Representative images of KPC mice with low and high levels of cfDNA. **d,** Quantification of cfDNA and **e,** representative images of KPC mice originating from another animal house (CRCM Marseille). N is indicated.

**Supplementary Figure 11: Description of the KPC cohort. a,** Enrolment criteria for KPC mice included in the study. **b,** Detailed description of KPC study mice.

**Supplementary Figure 12: Tumour progression by US imaging.** Representative US images of tumour progression in KPC mice treated with **a,** Vehicle and with **b,** BYL-719.**c,** Summary of tumour progression in the treated KPC mice.

**Supplementary Figure 13: Effects of PI3Kα pharmacological inhibition on cfDNA and metastasis. a,** Significant correlation between cfDNA and cancer cell proliferation in treated KPC mice. **b,** Representative images of (H&E and Ki67 IHC) metastases in liver, lung and spleen of KPC mice treated with the vehicle or BYL-719. N is indicated. Mean +/− SEM (* p<0.05, ** p<0.005, *** p<0.0001).

**Supplementary Figure 14: Cancer staging and tumour histology of KPC mice treated with the vehicle and BYL-719. a,** Cancer staging of KPC mice treated with the vehicle and BYL-719. **b,** Representative images of tumours in KPC mice. H&E, Movat’s Pentachrome and Picro Sirius Red staining.

**Supplementary Figure 15: Secondary effects of PI3Kα pharmacological inhibition. a,** Insulin quantification on treated KPC. **b,** Quantification of weight change in treated KPC mice. **c,** Representative images and quantification of cleaved Caspase 3 and **d,** γH2AX–positive cells in pancreatic sections of treated KPC mice. Mean +/−SEM (* p<0.05, ** p<0.005, *** p<0.0001)

**Supplementary Figure 16: Pharmacological inactivation of PI3Kα reverses pre-cancer lesions and alterations of their stromal environment in an oncogenic KRAS context.** a. Dosage regimen of PI3Kα-targeting drug (GDC0326 10mg/kg daily-blue arrows) in KC mice subjected to acute pancreatitis protocol with caerulein as described in Baer et al (black arrows). b-f IHC, stainings and quantification as indicated after 17 days; n=4 mice in each group. The surface of the lesion areas was quantified on two whole slides for each mouse, and normalised with the total area of the pancreas. Values are mean ± SEM. T-Test: *=P<0.05 Representative images are shown. A: acinar cells, E: epithelial lesion; FL=fibroblasts, S: stroma, I: Langerhans islet

**Supplementary Figure 17: Genotyping of murine cell lines derived for the study.**

**Supplementary Figure 18: Effects of PI3Kα inhibition on the immune response. a,** Representative images and quantification of CD4 and CD8 positive cells in KPC mice treated with the vehicle or BYL-719. N is indicated. **b,** Representative images of F4/80 and CD206 positive cells in KC and oncogenic PI3Kα mice at different ages. **c,** Representative images of CD206 positive cells around low-grade PanIN lesions in KC; p110α^+/+^, KC; p110α^+/lox^ and KC; p110α^lox/lox^ mice (4.5 months). **d,** Representative images of CD206 positive cells around high-grade PanIN lesions in KC; p110α^+/+^ and KC; p110α^+/lox^ (>6 months). **e,** Representative images of CD206 positive cells in KPC mice with normal, high-grade PanIN, PDACloc and PDACmet pancreas.

**Supplementary Table 1**: Collapsed gene expression in pancreatic samples from 36 normal, primary tumours from metastatic (PDACmet), localised pancreatic adenocarcinoma (PDACloc), chronic pancreatitis (CP) patients. Refers to Figure 1a-c. **a,** Hallmark scoring in normal, primary tumour from metastatic (PDACmet) or localised pancreatic adenocarcinoma (PDACloc), chronic pancreatitis (CP). Refers to Figure 1a. **b,** Reactome scoring in normal, primary tumour from metastatic (mPDAC) or localised pancreatic adenocarcinoma (IPDAC), chronic pancreatitis (CP). Refers to Figure 1b. **c,** PI3Kα gene signature scoring in normal, primary tumour from metastatic or localised pancreatic adenocarcinoma, chronic pancreatitis. Refers to Figure 1c.

**Supplementary Table 2: a,** PI3Kα gene signature scoring in TGCA. Refers to Figure 1d. **b,** PI3Kα gene signature scoring in PACA-AU. Refers to Figure 1d. **c,** PI3Kα gene signature scoring in GSE21501. Refers to Figure 1e.

**Supplementary Table 3:** Distribution of genetic alterations in PAAD cohort.

**Supplementary Table 4:**
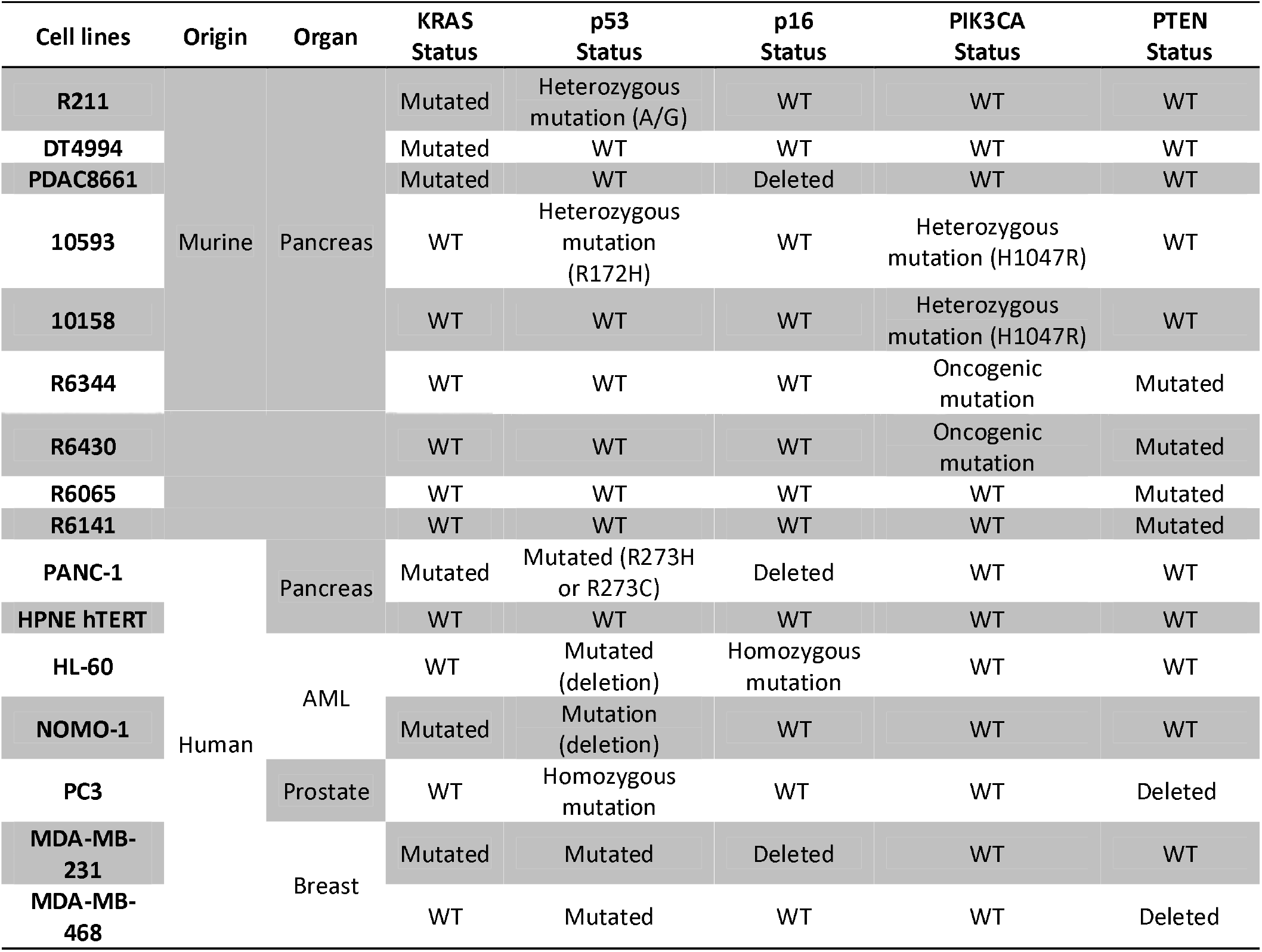
Information on cell lines. All cells were cultured in DMEM (Dulbecco’s Modified Eagle Medium), 4.5 g of glucose supplemented with 10% foetal bovine serum (FBS), 1% L-glutamine, 1% penicillin/streptomycin and 0.01% plasmocin, except for HL-60, NOMO-1 and PC3 cultured in RPMI (Roswell Park Memorial Institute) supplemented as described previously, and HPNE hTERT and HPDE cultured in 75% DMEM without glucose/25% medium M3 Base supplemented with 5% FBS, 10 ng/ml human recombinant EGF, 5.5 mM D-glucose (1g/L) and 750 ng/ml puromycin. Cells were cultured at 37°C in a humidified 5% CO_2_ atmosphere. AML: Acute myeloid leukaemia; WT: Wild type. R211-Luc cells were gifted by Pierre Cordelier and are R211 cells modified to express the luciferase. Cells were checked monthly by PCR for mycoplasma infection using the following primers: Forward: GCTGTGTGCCTAATACATGCAT and Reverse: ACCATCTGTCACTCTGTTAACCTC. Genomic DNA from a mycoplasma infected cell line was used as a positive control. Cell line authentication was presented in supplementary table 9.

**Supplementary Table 5:** KPC mice supplemental information, including cohort size for each experiment: refers to Figures 4 and 5. Power test analyses for KPC mice: refers to Figures 4 5.

**Supplementary Table 6:** Immune cell signatures scoring in PAAD cohort.

**Supplementary Table 7:** List of antibodies used for WB.

| Primary Antibody | Size (kDa) | Origin | Dilution | Company | Secondary Antibody |
| --- | --- | --- | --- | --- | --- |
| **p-Akt S473** | 60 | Rabbit | 1:1000 | Cell Signaling #4060 | 1:3300 |
| **p-Akt T308** | 60 | Rabbit | 1:1000 | Cell Signaling #9275 | 1:3300 |
| **Total Akt** | 60 | Rabbit | 1:1000 | Cell Signaling #4691 | 1:20000 |
| **Beta Actin** | 42 | Mouse | 1:1000 | Sigma Aldrich AC74 | 1:10000 |

**Supplementary Table 8:**
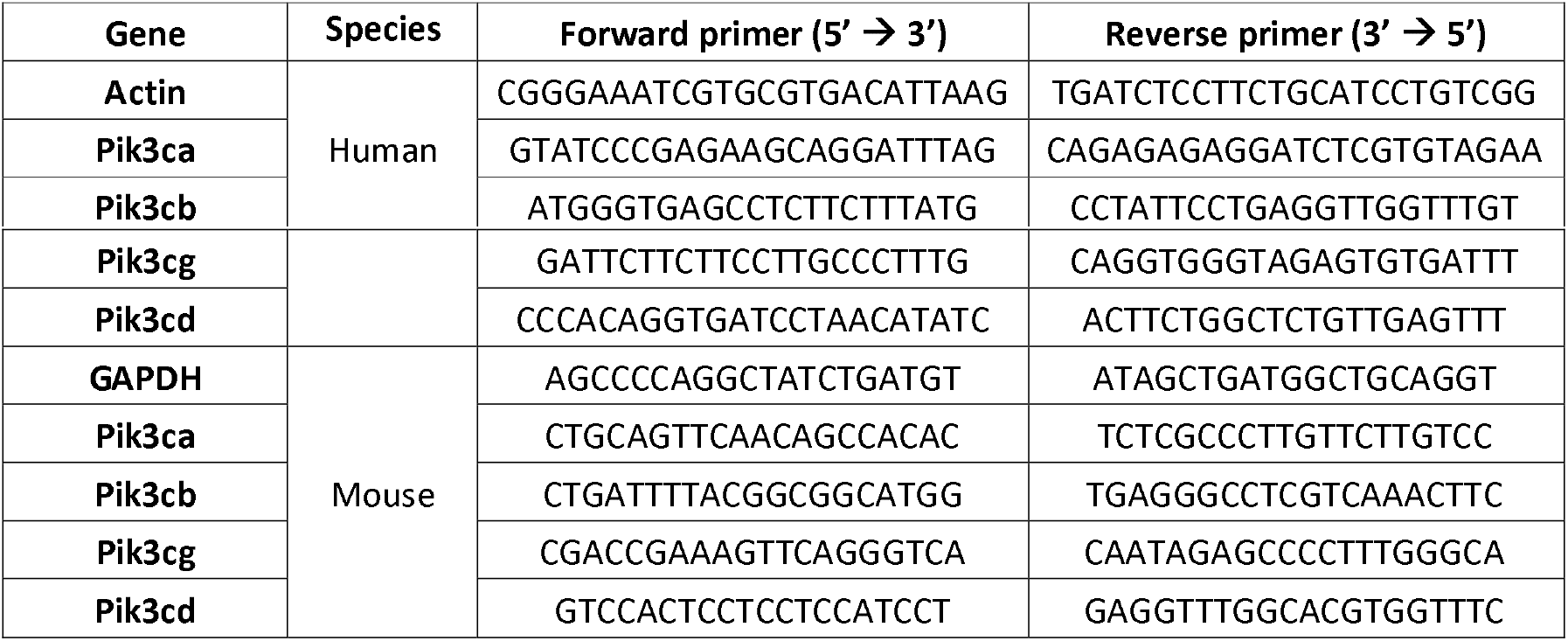
List of primers for qPCR.

**Supplementary Table 9:** Cell line authentication. Cell lines were authenticated by Eurofins Genomics (Germany). Genetic characteristics were determined by PCR-single-locus-technology. 16 independent PCR-systems D8S1179, D21S11, D7S820, CSF1PO, D3S1358, TH01, D13S317, D16S539, D2S1338, AMEL, D5S818, FGA, D19S433, vWA, TPOX and D18S51 were investigated. In parallel, positive and negative controls were carried out yielding correct results. Short Tandem Repeat (STR) similarity search was done with CLASTR 1.4.4 (https://web.expasy.org/cellosaurus-str-search/).

## Supplementary methods

### Human pancreatic samples

Patient samples from BACAP collection were collected after obtaining informed consent in accordance with the Declaration of Helsinki and stored with the “CRB Cancer des Hôpitaux de Toulouse”. According to French law, the BACAP collection has been declared to the Ministry of Higher Education and Research (DC-2008-463) and transfer agreements (AC-2008-820) (AC-2013-1955) have been obtained following approval by ethical Committees. Clinical and biological annotations of the samples have been declared to the CNIL (Comité National Informatique et Libertés – French national Data Protection Agency). All patient records and information were anonymized and encrypted prior to analysis.

### Transcriptomics and bioinformatics analysis

We selected transcriptional profiling datasets of normal pancreas, chronic pancreatitis and pancreatic primary tumoural tissues from localised (PDACloc) or metastatic patients (PDACmet). Published data on human samples were retrieved from public databases E_EMBL_6 (52) from compatible platforms, normalised using the RMA method (R 3.2.3, bioconductor version 3.2), collapsed (collapse microarray), filtered (SD>0.25) and statistically tested using an ANOVA test corrected using the Benjamini & Hochberg method (BH). For each sample, individual scoring for hallmarks or Reactome (actualised list of genes downloaded from MSigDB version [software.broadinstitute.org/gsea/msigdb] and Reactome [www.reactome.org]) was performed using Autocompare_SES software (available at https://sites.google.com/site/fredsoftwares/products/autocompare_ses) using the “greater” (indicating an enriched gene set) Wilcoxon tests with frequency-corrected null hypotheses (53), followed by values in each group of patients compared using an ANOVA test. Hierarchical patient clustering was performed using the PI3Kα activation signature. The PI3Kα gene signature was designed as the intersection of genes up-regulated and down-regulated in 20 breast tumours after BYL-719 treatment (54) and LINCS shRNA CMAP sig gene list (55)(suppl. Fig. 1). This list was narrowed down to 20 genes, which expression was found compatible with the quality criteria (filter) detailed above. Unsupervised hierarchical clustering of the E_EMBL_6 data set was performed focusing on the expression of these 20 genes regulated by PI3Kα in cancer.

Confirmed PDAC samples from public data bases were selected for further bioinformatics analysis. In detail, mRNA expression data and clinical data from confirmed PDAC patients of PAAD (TCGA) (175 patients) and PACA-AU (267 patients) cohorts were retrieved. Amongst the 175 well annotated TGCA patients, 21 patients were considered localised according to their UICC staging (T=0, 1 or 2, N=0, M=0). For each patient, a PI3Kα activation signature score or immune cell infiltration score (to quantify LTγδ, NK, LT CD8, Monocyte-Macrophage-DC, B cells, granulocytes, LT CD4) was given using SES auto compare software, and patients were hierarchically clustered in 3 groups corresponding to high, medium or low scoring. Scores in each group were statistically tested using an ANOVA test corrected according to the Benjamini & Hochberg method (BH). For PI3Kα activation signature, high and medium groups were then pooled. The overall survival of patients in each cluster was plotted and statistical differences were calculated using the log rank test. The prognosticator value of PI3Kα activation signature and IUCC staging were tested independently and compared to the value of clinical T,N,M staging using the multivariate Cox test and the PAAD data base. We verified that within each patient cluster, there was no enrichment in terms of genetic changes associated with the PI3K/Akt pathway and, in particular, that oncogenic mutations of PI3Kα and PTEN were infrequent and equally distributed in each group of patients (mutational pattern available in PAAD cohort only: suppl. Table 3, suppl. Fig. 2).

### Mice

The LSL-KRAS^G12D^ and LSL-p53^R172H^ knock-in (from D. Tuveson, Mouse Models of Human Cancers Consortium Repository, Frederick National Cancer Institute), Pdx1-Cre (from D.A. Melton, Harvard University, Cambridge, MA), Pdx-1Cre (from D. Tuveson) and p110α^lox/lox^ (from B. Vanhaesebroeck, University College London) strains were interbred against a mixed background (CD1/SV129/C57Bl6) to obtain the compound mutants LSL-Kras^G12D^-LSL-p53^R172H^-Pdx1-Cre (KPC), LSL-KRAS^G12D^-Pdx1-Cre (KC), LSL-KRAS^G12D^-Pdx1-Cre-p110α^+/lox^ (KC;p110α^+/lox^), KRAS^G12D^-Pdx1-Cre-p110α^+/lox^ (KC; p110α^lox/lox^), LSL-KRAS^G12D^-LSL-p53^R172H^-Pdx1-Cre-p110α^+/lox^ (KPC; p110α^+/lox^), LSL-KRAS^G12D^-LSL-p53^R172H^-Pdx1-Cre-p110α^+/lox^ (KPC; p110α^lox/lox^). Littermates not expressing Cre were used as controls. KPC mice were bred in three animal houses (mixed background, Melton’s Pdx1-Cre: CRCT, Anexplo, Toulouse, France, 2 sites) and (C57B6 background, Tuveson’s Pdx1-cre: CRCM, Marseille, France). The Ptf1a^Cre/+^-LSL-KRAS^G12D/+^ (KC; p110α^+/+^) and Ptfla^Cre/+^-LSL-PIK3CA^H1047R/+^ (p110α^H1047R^) strains were bred at the Technische Universität München (Technical University, Munich). Genotyping was performed as described in (9, 24, 56) and analysed with the Fragment Analyzer Systems (AATI) DNA Analysis Kits dsDNA 910 Reagent kit, 35-1500bp (AATI). The genotyping of p53 allele uses T035-5’-CTTGGAGACATAGCCACACTACTG-3’/ T036 - 5’-AGCTAGCCACCATGGCTTGAGTAAGTCTGCA-3’ / T037 - 5’-TTACACATCCAGCCTCTGTGG-3’-Annealing temperature: 60°C – amplicon size for WT: 166bp, for LSL-p53R172H: 270bp. Animal sample size was calculated using a power analysis in order to obtain statistically significant results from ki67 (n=5, p≤0.02) and CD206 (n=6, p≤0.05) immune staining (suppl. Table 5).

Mutant KRAS, with or without partially deficient PI3Kα activity pancreatic cancer cell lines were derived from the pancreas and lung of KC or KC;p110α^+/lox^ animals (aged 10-13 months): organs were dissected using a scalpel in DMEM containing 200 U/ml of collagenase (Sigma C6885), incubated for 20 minutes in a water bath at 37°c under agitation. After washing, the cell suspension was resuspended in 10% DMEM FBS, filtered through a 100 μm sieve and seeded in a 10-cm cell dish. After 7 passages, genotyping was carried out (suppl. Fig. 17) with confirmed recombination of KRAS (PCR for detection of unrec (positive control: R211 cell line). LSL-KRASG12D allele: K005 5’-AGCTAGCCACCATGGCTTGAGTAAGTCTGCA-3’ / K008 5’-TCCGAATTCAGTGACTACAGATGTACAGAG-3’ – annealing temperature: 55°C - size of amplicon: 300bp; PCR for detection of recombined LSL-KRASG12D: K001 5’-GTCTTTCCCCAGCACAGTGC-3’ / K002: 5’-CTCTTGCCTACGCCACCAGCTC-3’ - annealing temperature: 61°C - size of amplicon: doublet at 554bp) without (R211, A338, A338L cell lines) or with *Pik3ca* gene recombination (A260, A94L cell lines) (PCR for detection of loxed Pik3cα gene: FE1-5’-GGATGCGGTCTTTATTGTC-3’ / FE4-5’-TGGCATGCTGCCGAATTG-3’ - annealing temperature: 59°C - amplicon size for WT: 640bp, for unrec. lox: 708 bp; PCR for detection of rec *Pik3ca* gene: aRecF1 5’-GGGGACAGTAGGAGGATGGT-3’ / aRecR 5’-TGGCATTCCAGAGCCAAGC-3’ - annealing temperature: 65°C - amplicon size: 812bp); it should be noted that recombination is driven by Cre expression in pancreatic lineage (Pdx1-cre: H06: 5’-TGCCACGACCAAGTGACAGCAATG-3’; H07 5’-GACCAGAGTCATCCTTAGCGCCG-3’ – annealing temperature: 65°C - amplicon size: 580bp). Tumorigenicity of A338 and A260 cell lines was verified by injection (7.5×10^4^ or 1.25×10^5^ cells) in Nude and NSG mice. Cells were passaged several times before experiments to avoid stromal cell contamination.

Pancreatic injury was induced on young KC mice (8-12 weeks old) by a series of six hourly intraperitoneal injections of caerulein (75 μg/kg of body weight, Bachem) repeated after 48 h. Eight days later, a treatment/intervention protocol was developed to allow formation of pre-cancer lesions: mice were treated with the vehicle (0.5% methyl cellulose with 0.2% Tween-80) or with GDC0326 (10mg/kg) by gavage (suppl. Fig. 16a).

### Cell culture

All the cell lines described in supplementary Table 4 were obtained from the American Type Culture Collection (ATCC, Manassas, VA) or from genetically engineered mouse models or from CRB, Toulouse, IUCT-O. All cells were cultured in DMEM (Dulbecco’s Modified Eagle Medium), 4.5 g of glucose supplemented with 10% foetal bovine serum (FBS), 1% L-glutamine, 1% penicillin/streptomycin and 0.01% plasmocin, except for HL-60, NOMO-1 and PC3 cultured in RPMI (Roswell Park Memorial Institute) supplemented as described previously or HPNE hTERT that were cultured in 75% DMEM without glucose/25% medium M3 Base supplemented with 5% FBS, 10 ng/ml human recombinant EGF, 5.5 mM D-glucose (1g/L) and 750 ng/ml puromycin. Cells were cultured at 37°C in a humidified 5% CO_2_ atmosphere. AML: Acute myeloid leukaemia; WT: Wild type. R211-Luc cells are R211 cells modified to express the luciferase. Cells were checked monthly by PCR for mycoplasma infection using the following primers: Forward: GCTGTGTGCCTAATACATGCAT and Reverse: ACCATCTGTCACTCTGTTAACCTC. Genomic DNA from a mycoplasma-infected cell line was used as a positive control.

### Inhibitors

All PI3K inhibitors for in vitro use were purchased (CliniSciences) and dissolved in dimethyl sulfoxide (DMSO) to obtain a stock concentration of 10 mM, subsequently diluted as indicated and compared to the diluted DMSO vehicle (vehicle). In vivo, BYL-719 (ApexBio) was dissolved in 0.5% methyl cellulose with 0.2% Tween-80 and administered by oral gavage at 50mg/kg daily.

Inhibitory concentration 50 (IC50) in nM identified in vitro on recombinant proteins of all PI3K inhibitors tested for p110α, p110β, p110δ, p110γ and mTOR. NA: not available.

| Inhibitor | p110α IC<sub>50</sub><br>(nM) | p110β IC<sub>50</sub><br>(nM) | p110δ IC<sub>50</sub><br>(nM) | p110γ IC<sub>50</sub><br>(nM) | mTOR<br>(nM) |
| --- | --- | --- | --- | --- | --- |
| **A66** (57) | **32** | > 12500 | > 1250 | 3480 | > 5000 |
| **BYL-719** (58) | **4.6** | 1156 | 290 | 250 | > 9100 |
| **GDC-0326** | **0.2** | 133 | >1250 | 3480 | >5000 |
| **TGX-221** (48) | 5000 | **5** | 100 | > 10000 | NA |
| **TGX-155** (60) | > 20000 | **30** | 340 | > 20000 | NA |
| **TGX-115** (61) | 61000 | **130** | **630** | 100000 | > 100000 |
| **AZD8186** (62) | 35 | **4** | **12** | 675 | NA |
| **AS252424** (63) | 940 | 20000 | 20000 | **30** | NA |
| **LY294002** (63) | 720 | 306 | 1330 | 7260 | NA |
| **BKM120** (64) | 52 | 166 | 116 | 262 | 4600 |
| **GDC0941** (65) | 3 | 33 | 3 | 75 | 580 |
| **PI-103** (66) | 2 | 3 | 3 | 15 | 30 |

### Cytotoxicity assay

Cells were seeded in 96-well plates. Twenty-four hours later, they were treated with a dose range of α-selective, β-selective, β/δ-selective, γ-selective or pan PI3K inhibitors at 0.1, 1 and 10 μM, respectively, in 10% FBS medium or the vehicle. Three days post-dose, living cells were stained using MTT for adherent cell lines or MTS for acute myeloid cell lines. Control cells were treated with 0.1% DMSO and considered as 100% living cells. Each condition was conducted in triplicate and each experiment was performed at least three times.

### Motility assay

Cells (1×10^5^) were seeded in 24-well plates. Scratching was carried out at confluence using a pipette tip and cells were treated with α-selective and pan-PI3K inhibitors at 0.01, 0.1 or 1 μM, or untreated (vehicle). Cells were examined microscopically after 8 h and 24 h, and the scratch surface was analysed using ImageJ software. Control cells were treated with 0.01% DMSO. Five photographs were taken per condition and each experiment was carried out at least three times.

### Migration assay

Cells (5×10^4^) were seeded in the upper compartment of Boyden chamber inserts in serum-free DMEM. Four hours after seeding, complete DMEM was added to the lower compartment to trigger migration. Cells were treated with α-selective, β-selective, β/δ-selective, γ-selective and pan-PI3K inhibitors at 0.1 and 1 μM or the vehicle in the lower and upper compartments. Twenty-four hours after treatment, cells were fixed with 3.7% formaldehyde for 20 minutes and then stained with crystal violet for 20 minutes. A cotton swab was used to remove non-migrated cells from the insert. Migrated cells were analysed using ImageJ software. Control cells were treated with 0.01% DMSO. Ten photographs were taken per condition and each experiment was carried out at least three times.

### Western blot

Cells were scraped in cold lysis buffer (50mM Tris at pH 7.4, 150 mM NaCl, 1% Triton, 1 mM EDTA, 2 mM DTT, 2 mM NaF, 4 mM Na orthovanadate and a protease inhibitor cocktail provided by Roche). Fifty micrograms of proteins were separated by SDS-PAGE and transferred to a PVDF membrane. Membranes were saturated in TBS (50 mM Tris, 150 mM NaCl)/0.1% Tween 20 (TBST)/5% milk and incubated with the primary antibody (suppl. Table 7). Membranes were washed with TBST and incubated with the corresponding secondary antibody. Membranes were washed with TBST and immunocomplexes were visualised using ECL RevelBlot Plus (Ozyme) by autoradiography.

### siRNA

Cells were transfected with Lipofectamine 2000 (ThermoFisher Scientific) with SMARTpool ON-TARGETplus mouse siRNA (Dharmacon) targeting: Pik3ca, Pik3cb, Pik3cg, Pik3cd according to the manufacturer’s protocols as used in (67–69). ON-TARGETplus Non-targeting control siRNAs (Dharmacon) were used as controls. Twenty-four hours after transfection, the cells were used for migration or RT-qPCR experiments.

### RNA extraction and RT-qPCR

The cell pellet was resuspended in Trizol (Invitrogen). Chloroform was added and the suspension was centrifuged for 10 minutes at 18,000 g at 4°C. The aqueous phase was isolated, completed with isopropanol then centrifuged for 10 minutes at 18,000 g at 4°C. The pellet was washed with cold 80%, 83% EtOH. The suspension was centrifuged for 5 minutes at 18,000 g at 4°C and the pellet was dried and then resuspended in RNAse-free water. 1 μg of extracted RNA was used to obtain cDNA using RevertAid H minus reverse transcriptase, random hexamers and the corresponding mix (ThermoScientific). Primers were all designed with Primer-BLAST (NCBI) and are listed in supplementary Table 8. qPCRs were produced using the SsoFast EvaGreen supermix (Bio-Rad). Expression was normalised with Actin primers.

### Tail vein injection in Nude mice

5×10^4^ R211-Luc cells were injected into the tail vein of Nude mice. The mice were treated 5 days a week for 3 weeks starting on the injection date. Two and 3 weeks post-injection, mice were injected i.p. with 150 mg/kg of RediJect D-Luciferin (Perkin Elmer) and monitored for Luciferase expression after 6-8 minutes in the IVIS Spectrum in vivo imaging system (Perkin Elmer). Luminescence (photo count) was measured for each mouse.

### Tumour detection by ultrasound imaging

Ultrasound imaging was performed using the VisualSonics Vevo2100 High Resolution System equipped with an ultrasound transducer in the 25-55MHz range. Animal preparation and imaging procedures were performed as described in Sastra *et al* (70). KPC mice were monitored once a week from 12 weeks old onwards; when a tumour was detected, ultrasound scans were performed every other day. The tumour area was measured by delimiting the tumour border and determining the major axis; at least 5 replicates were performed per mice per ultrasound. Tumour volume was calculated using the formula V= (4/3) x *π* x (Length/2)^2^ x (Depth/2). The tumour volume fold change corresponds to the increased fold change in the tumour volume after treatment onset.

### Blood collection and plasma separation

Blood samples were collected every two weeks via retro-orbital collection from the age of 12 weeks onwards. A maximum volume of 100 – 150 μL was collected on each occasion. The blood was collected using Pasteur glass pipettes and then transferred to Eppendorf tubes containing 20 μl 0.5M EDTA. Plasma was separated by centrifuging blood at 1500 g for 20 min at 4°C within 3h of blood collection. The plasma was stored at −80°C until required for further use.

### cfDNA extraction from blood plasma

For cfDNA extraction, blood plasma was re-centrifuged at 18,000 g for 10 min at room temperature to reduce debris contamination. cfDNA was extracted from blood plasma using the QIAmp DNA Mini Kit (QIAGEN) protocol except for eluting the cfDNA in 50μL of elution buffer. cfDNA samples were stored at −20°C until required for further use. CfDNA quantification was performed by qPCR as described in the supplementary methods.

### Blood count

Blood counts were performed using Yumizen H500 hematology analyzer (HORIBA), calibrated for murine blood.

### Quantification and characterisation of cfDNA

For quantifying the relative cfDNA in blood plasma, we initially plotted a standard curve using extracted DNA from a murine cell line R211 (without LSL cassette, expressing mutated *KRAS* and *TP53*) and from pancreatic extracts from mice expressing the LSL cassette (not recombined). The qPCR was performed using 1 μL of DNA and SsoFast Eva Green Supermix (BioRad^®^). For the standard curve, we did serial dilutions up to 1/1000 of the two DNA extracts and a qPCR of two different genes, p53 and GAPDH. We designed and selected the most specific and efficient primers (Sigma-Aldrich^®^) using Primer-Blast (NCBI). The standard curves thus obtained allowed subsequent quantification of cfDNA in mouse plasma samples.

The following primers were used:

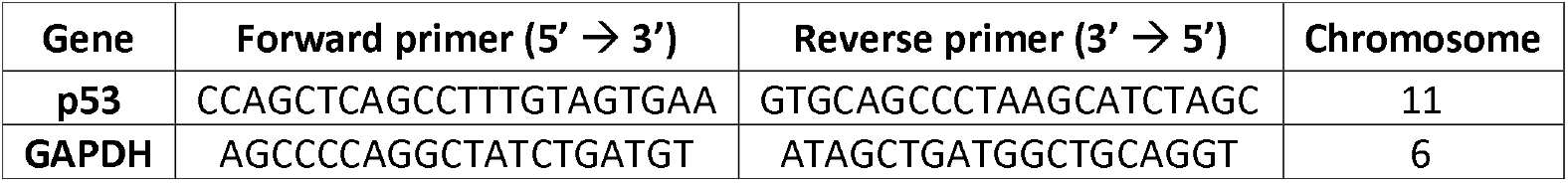

cfDNA thresholds were calculated by compiling all cfDNA measurements from mice with normal pancreas, high-grade PanINs, localised and metastatic PDAC. For the survival curve (Kaplan-Meier), mice were categorised as having a normal, medium or high level of cfDNA according to their mean cfDNA values. As for thresholds, the low range corresponds to mice presenting 0 – 0.003 AU of cfDNA and the high range to mice presenting cfDNA above the mentioned level. In order to establish these ranges, we analysed the histology of each mouse at end point. cfDNA was quantified using blood collected at sacrifice/death. All raw data and threshold calculations are shown in suppl. Table 5.

The Fragment Analyzer™ was used to determine the size of DNA fragments in blood plasma. The DNF-474 High Sensitivity NGS Fragment Kit was used to characterise cfDNA. Three different cfDNA fragmentation profiles were obtained: for normal and healthy mice, the electropherogram did not present any fragment; for mice with high-grade PanINs and localised PDAC, a 160-210bp fragment was always found; for mice with metastatic PDAC, the electropherogram presented the aforementioned 160-210bp fragment, in addition to larger fragments but at lower concentrations.

### Details of the IHC procedure and list of antibodies used

Tissues were fixed in 10% neutral-buffered formalin (Sigma HT501128) and embedded in paraffin. For pathological analysis, the tissues were serially sectioned (4 μm), and then stained with haematoxylin eosin. All tissues were analysed in blinded fashion. Histopathological scoring of pancreatic lesions was performed on sections 100 μm apart, with 3 sections per pancreas. IHC was performed and quantified as described in (9).

Immunostaining was conducted on formalin-fixed, paraffin-embedded tissues using standard methods. After rehydration, slides were permeabilised for 10 min in 0.1% Triton/1% PBS; they were then subjected to heat-antigen retrieval in citrate buffer or to enzymatic antigen retrieval in Proteinase K (Dako S3020), the latter was only applied for F4/80 immunostaining. Slides were incubated for 20 min with Protein Block (Dako X0909) to prevent non-specific binding. Sections were subsequently incubated overnight or for 1 h at room temperature* with primary antibodies (See Table below), washed, incubated for 15 min with 3% H_2_O_2_ and then rewashed. Most antibodies were detected using the HRP-Detection Reagent SignalStain^®^Boost (CST 8114) and developed through AEC (Dako K3464) incubation; anti-F4/80 antibody was revealed using the VECTASTAIN^®^ Elite^®^ ABC HRP Kit (VECTOR Laboratories PK-6100). All slides were counterstained with haematoxylin and mounted using Glycergel Mounting Medium (Dako C0563). Quantifications were randomly performed in at least 5 large-scale images in the area indicated.

| Primary Antibody | Origin | Dilution | Company |
| --- | --- | --- | --- |
| Anti-ki67 | Rabbit | 1:200 | Abcam<br>ab15580 |
| Anti-mannose receptor<br>(CD206) | Rabbit | 1:3000 | Abcam<br>ab64693 |
| Anti-CD4 | Rabbit | 1:100 | Abcam<br>ab183685 |
| Anti-CD8 | Rabbit | 1:400 | Abcam<br>ab203035 |
| Anti-cleaved Caspase 3 | Rabbit | 1:50 | Cell Signaling<br>#9664 |
| Anti-γH2Ax | Rabbit | 1:500 | Cell Signaling<br>#9718 |
| Anti-F4/80* | Rat | 1:100 | Invitrogen<br>MA1-91124 |
| Anti-phospho Akt<br>substrate | Rabbit | 1:1000 | Cell Signalling<br>#9611 |

We followed the manufacturers’ protocol for additional staining with the Picro Sirius Red Stain Kit (ab150681) and Movat’s Pentachrome Stain Kit (Modified Russell-Movat ab245884).

### Cytokine measurement (Luminex^®^ assay) in blood samples

A MILLIPLEX^®^_MAP_ (Merc Millipore # MCYTMAG70PMX32BK) assay was performed using 25 μL of non-diluted murine blood plasma.

We specifically tested for a panel of 32 murine chemokines and cytokines: Eotaxin, G-CSF, GM-CSF, IFNγ, IL-1α, IL-1β, IL-2, IL-3, IL-4, IL-5, IL-6, IL-7, IL-9, IL-10, IL-12 (p40), IL-12 (p70), IL-13, IL-15, IL-17, IP-10, KC, LIF, LIX, MCP-1, M-CSF, MIG, MIP-1α, MIP-1β, MIP-2, RANTES, TNFα, and VEGF. IL3 could not be detected on a comparative scale.

### Cytokine measurement in cell supernatants (Q-Plex™ assay)

The assay was performed according to the manufacturer’s protocol. A Q-Plex™ assay (Quansys Biosciences, Q-Plex mouse cytokine Stripwells 16-plex assay) was carried out to measure the concentration of 16 mouse cytokines in cell supernatants. The concentration was normalised with the number of living cells. R211, A338 and A260 were derived from primary tumours. A338L and A94L were isolated from lung metastases. All cell lines were passaged at least 15 times to avoid stromal contamination. KC;p110α^+/+^(R211, A338, A338L) and KC;p110α^+/lox^(A260, A94L) cells (3.10^6^) were seeded in 6-well plates with 10% FBS. On the following day, the medium was replaced by 2% FBS medium with or without the vehicle (0.01% DMSO) or 1 μM BYL-719. After 2 days of treatment, the medium was replaced by a new medium with the same content. The cell supernatant was extracted after one day.

### PIP extraction and measurement by mass spectrometry

More specifically, murine pancreatic tumour cells R211 were treated with or without the vehicle (0.01% DMSO), A66 1 μM or BKM120 1 μM for 15 minutes. The culture medium was quickly aspirated and cells were scraped in cold HC1 1M. The lysate was centrifuged for 5 minutes at 15,000 g at 4°C. The cell pellet was stored at −80°C until required for PIP extraction. Lipids were extracted and derivatised using TMS-diazo-methane as previously described (71). Mass spectral analysis was performed on the LC-QqQ triple quadrupole mass spectrometer (LC-vcQQQ 6460 Agilent) equipped with positive mode electrospray ionisation as described above (71). Analyses were performed in Selected Reaction Monitoring detection mode (SRM) using nitrogen as collision gas. Finally, peak detection, integration and quantitative analysis were performed using MassHunter QqQ Quantitative analysis software (Agilent Technologies VersionB.05.00) and Microsoft Excel software. Data were processed using QqQ Quantitative (vB.05.00) and Qualitative analysis software (vB.04.00).

### Measurement of cfDNA in Human patient plasma samples

Blood was collected from patients diagnosed with pancreatic adenocarcinoma and analysed by digital droplet PCR (ddPCR) for KRAS mutation. Briefly, 10 mL of blood were centrifuged twice in PAXgene blood ccfDNA tubes (Qiagen) at 1,200 g for 10 min at 4°C and 16,000 g for 10 min at 4°C. Cell-free plasma was collected and total DNA was extracted from 3ml of plasma using the QIAamp Circulating Nucleic Acid Kit (Qiagen) according to the manufacturer’s recommendation. Circulating cell-free DNA (cfDNA) ranging from 110-210 base pairs was qualified and quantified using the DNF-474 high sensitivity ngs fragment analysis kit (Agilent). For ddPCR, 5ng of cfDNA were analysed using the ddPCR™ KRAS G12/G13 Screening Kit #1863506 (Biorad) and the QX200 Droplet Digital PCR (ddPCR) System (Biorad), according to the manufacturer’s recommendation. Samples with high-molecular DNA were excluded from the analysis. Allelic frequency percentages for KRAS mutation were obtained using QuantoSoft software (Biorad).

### Patient and Public Involvement

This research was partly funded by the charity Fondation de France. Dissemination by BT was performed through Youtube: https://www.youtube.com/watch?v=GE_A-ApZ-cA. Both FRD and SA disseminated their research to high school students, as part of their commitment in MSCA-ITN funding.

### Statistical analysis

Experimental data provided at least 3 experimental replicates and five measurement replicates. Statistical analyses were performed with GraphPad Prism using T-tests (non-parametric Mann-Whitney): *p<0.05, **p<0.01, ***p<0.001. Correlation analysis was performed using Pearson r test. Statistical relevance of the cohort size was determined using a Power Calculation test (www.lasec.cuhk.edu.hk/.../power_calculator_14_may_2014.xls)

## Notes

Conflict of interest, EPT was funded by Cellgene.

### Competing Interest Statement

EPT received a Cellgene funding.

