## Supplementary material for "Pancreatic Cancer Intrinsic PI3Kα Activity accelerates Metastasis and rewires Macrophage Component": S1

**a**

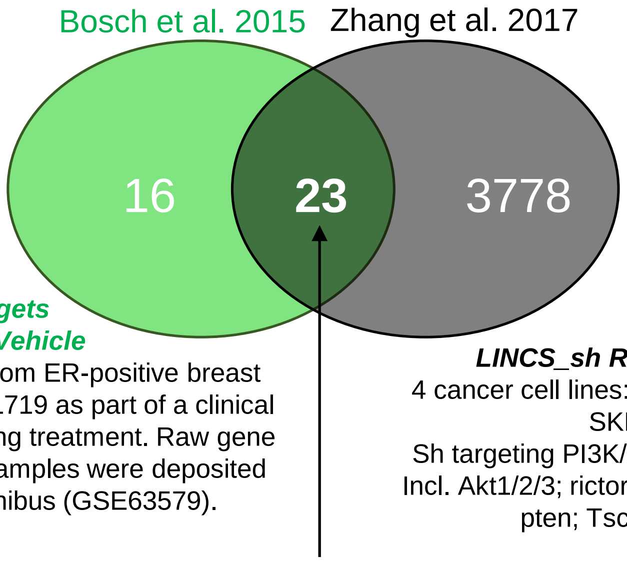


***PI3Kα targets***
***BYL-719 vs Vehicle***
20 Paired tumour biopsies from ER-positive breast cancer patients receiving BYL719 as part of a clinical trial collected before and during treatment. Raw gene expression data for patient samples were deposited in Gene Expression Omnibus (GSE63579).

***LINCS_sh RNA_CMAP sig***
4 cancer cell lines: PC3, HL60, MCF-7, SKMEL5
Sh targeting PI3K/AKT/MTOR pathway
Incl. Akt1/2/3; rictor; raptor; mtor; pik3ca; pten; Tsc1/2; Inpp4b

CCNB1, BIRC5, CCNE1, BCL2, PHGDH, ESR1, TYMS, MELK, FGFR4, GPR160, PTTG1, CDH3, MDM2, ERBB2, CEP55, EXO1, GRB7, MYBL2, KIF2C, CDC20, FOXA1, RRM2, SLC39A6

**Supplementary Figure 1**
