## Supplementary figures and images for "Pancreatic Cancer Intrinsic PI3Kα Activity accelerates Metastasis and rewires Macrophage Component"

### S2

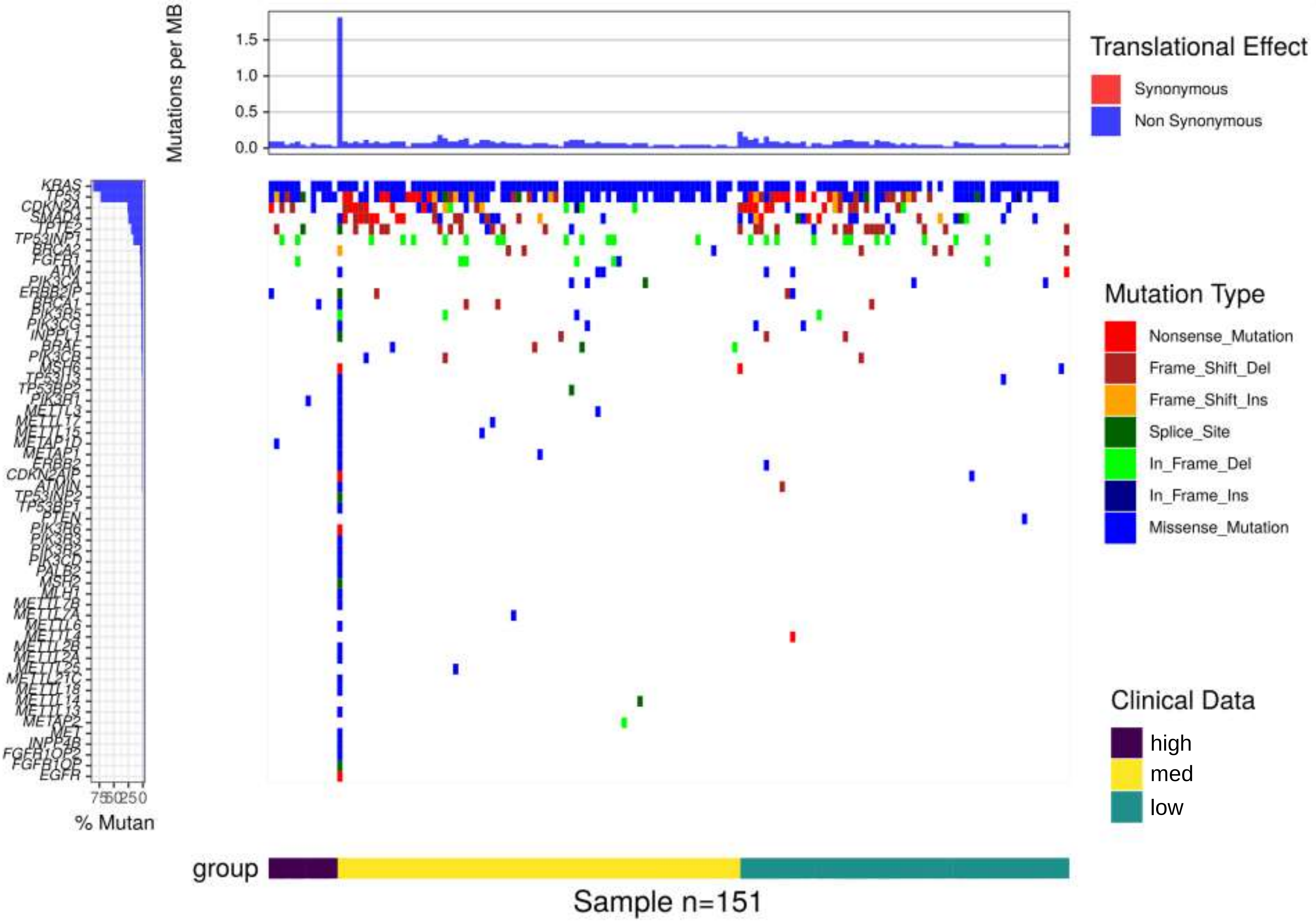


**Supplementary Figure 2**

### S3

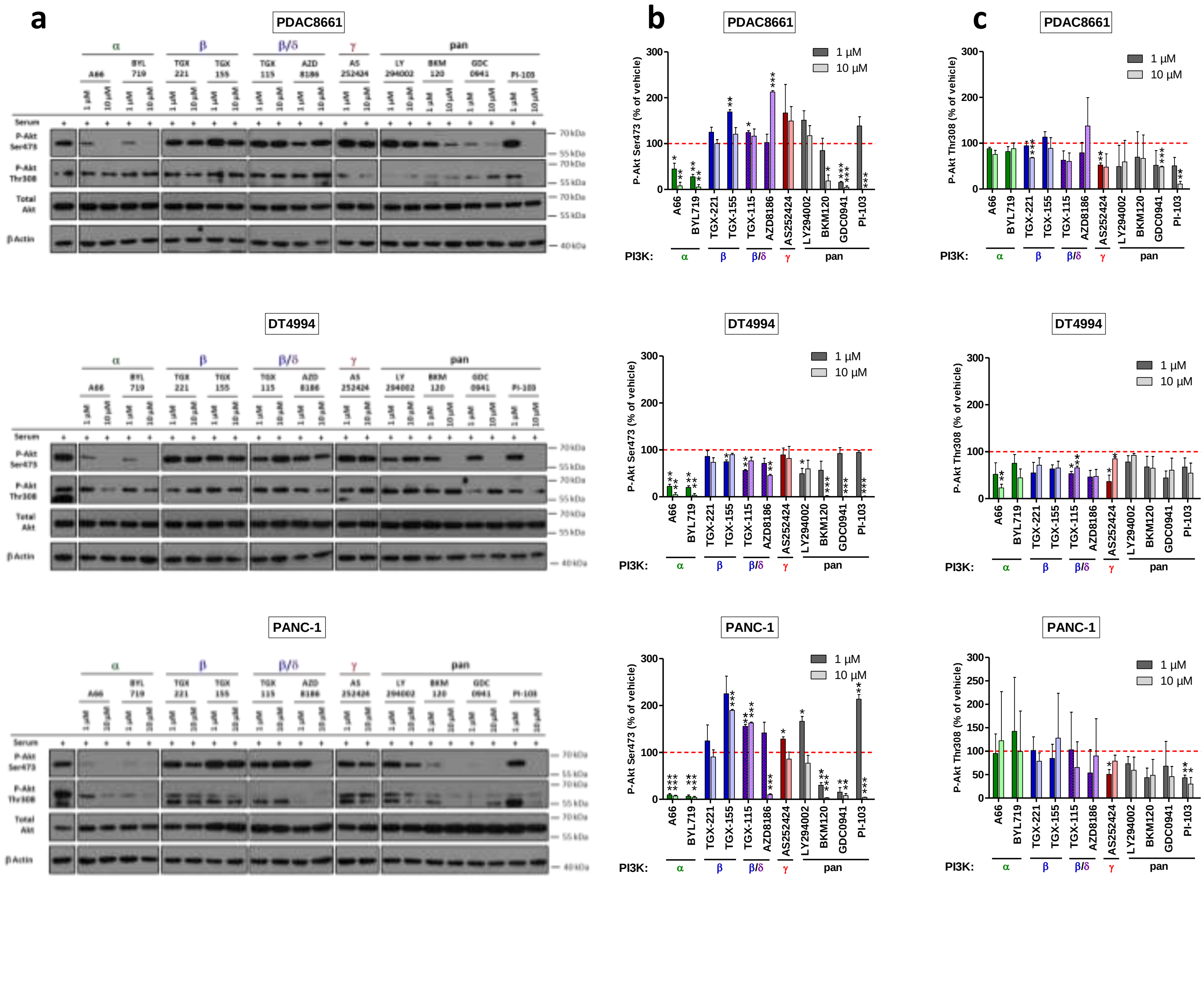


#### Supplementary Figure 3

### S4

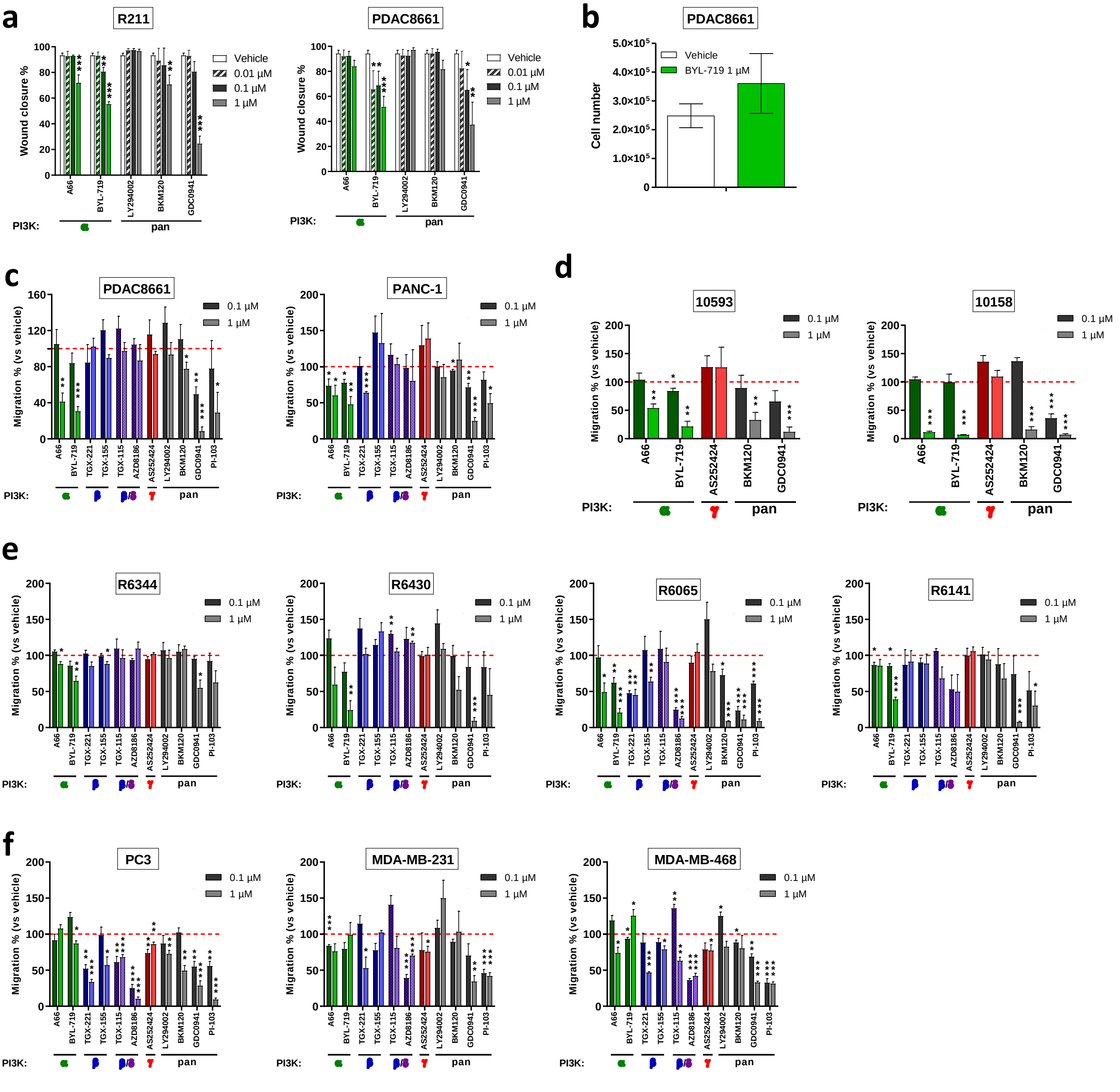


#### Supplementary Figure 4

### S5

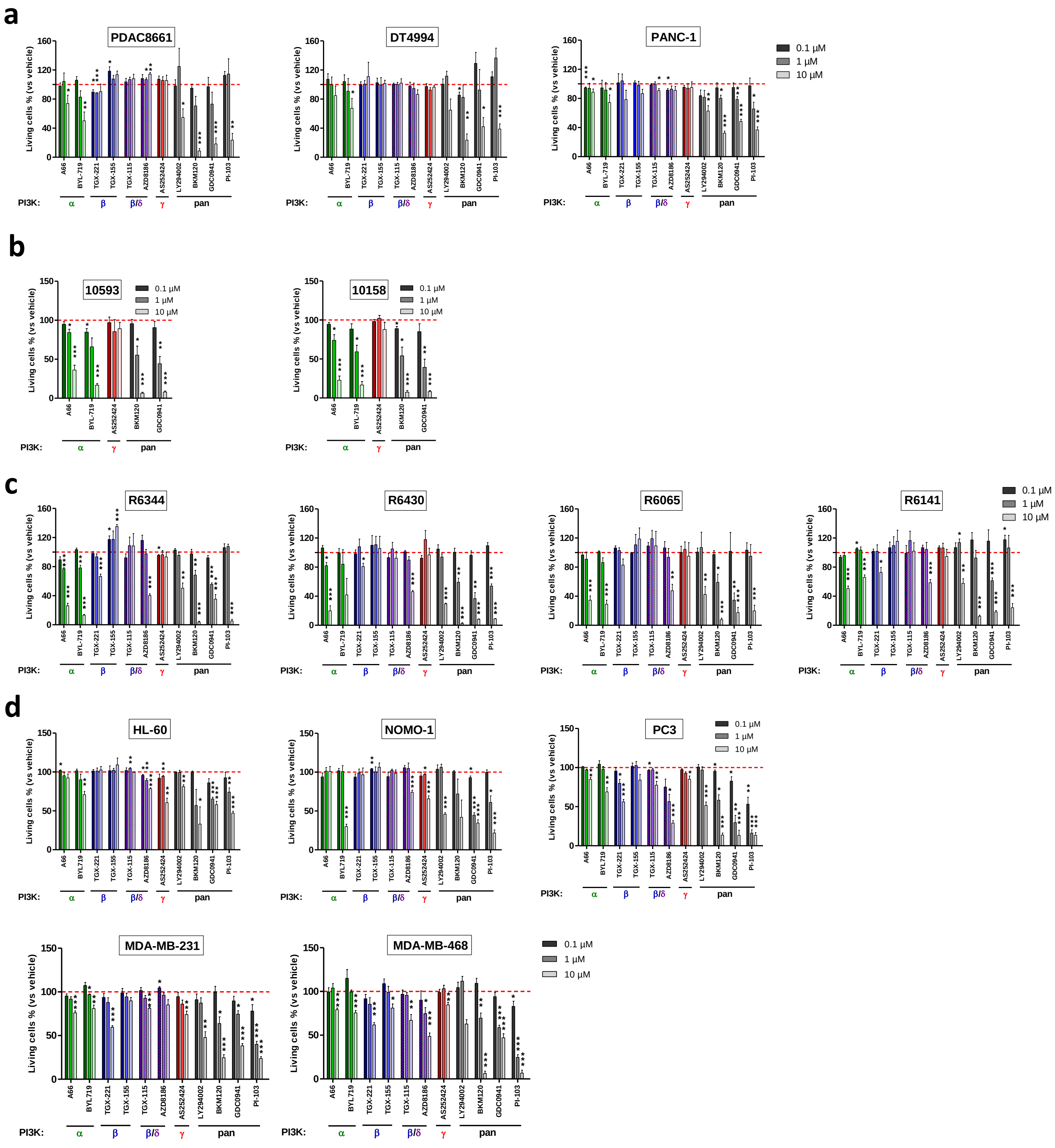


#### Supplementary Figure 5

### S7

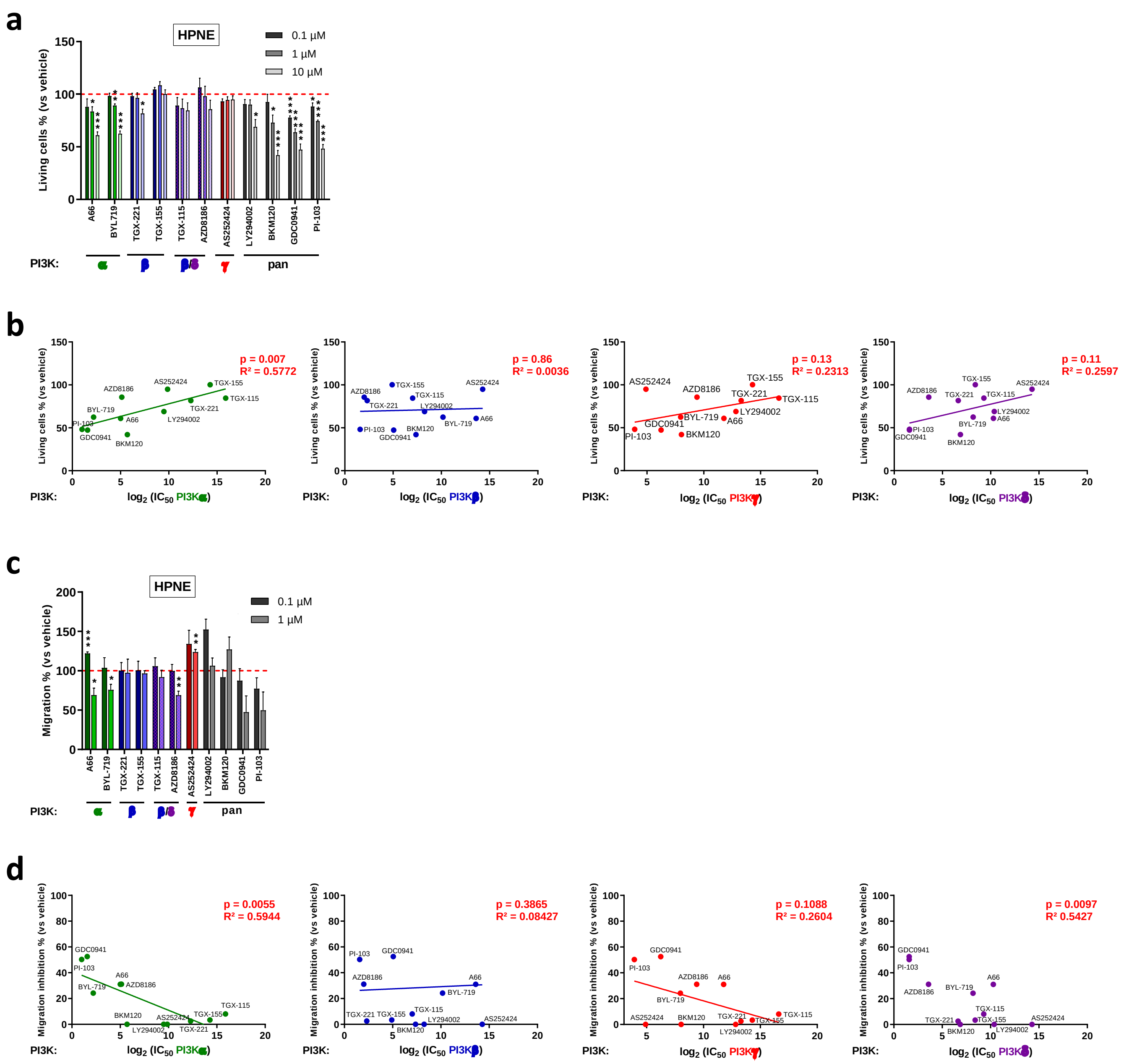


#### Supplementary Figure 7

### S8

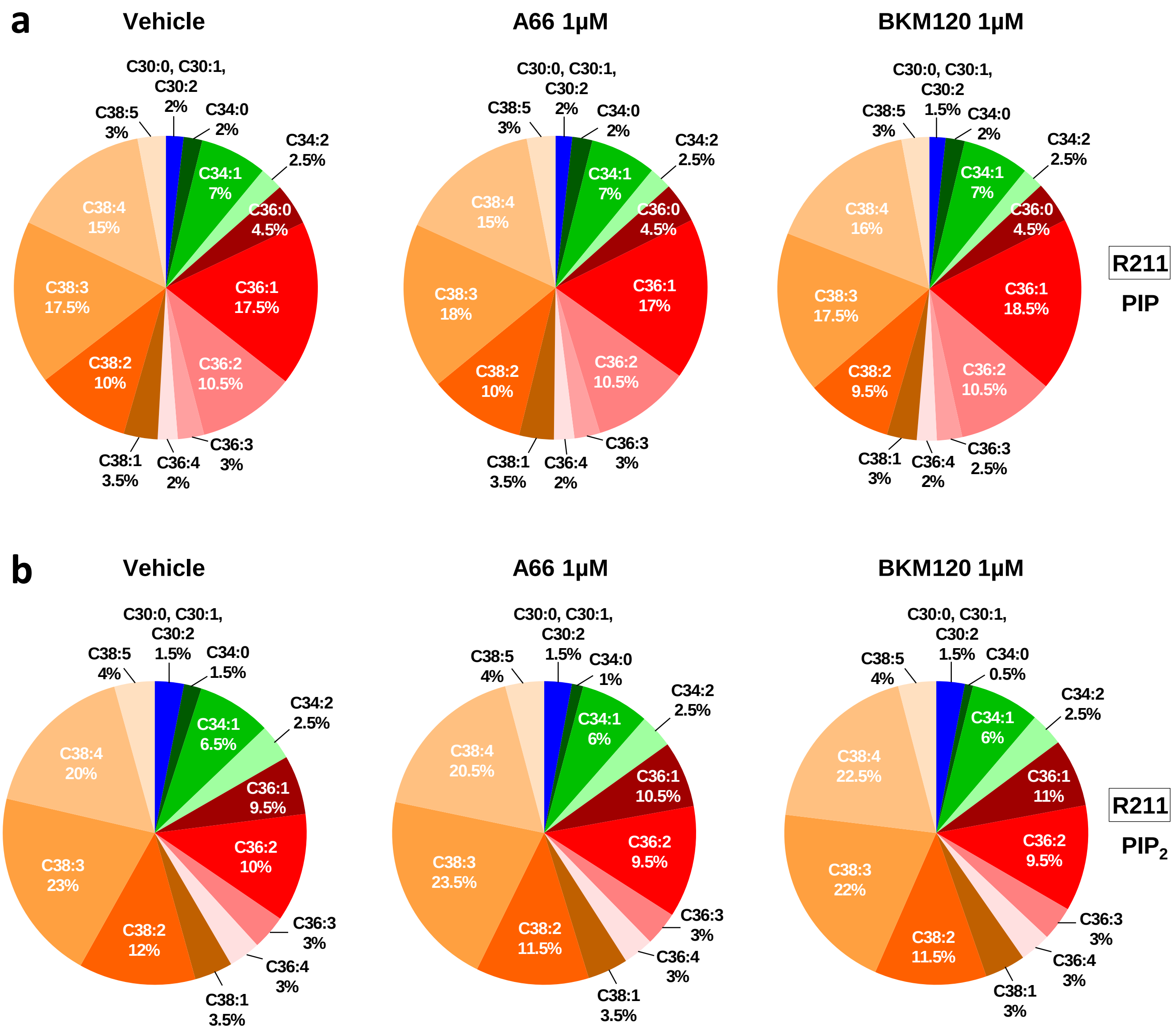


#### Supplementary Figure 8

### S10

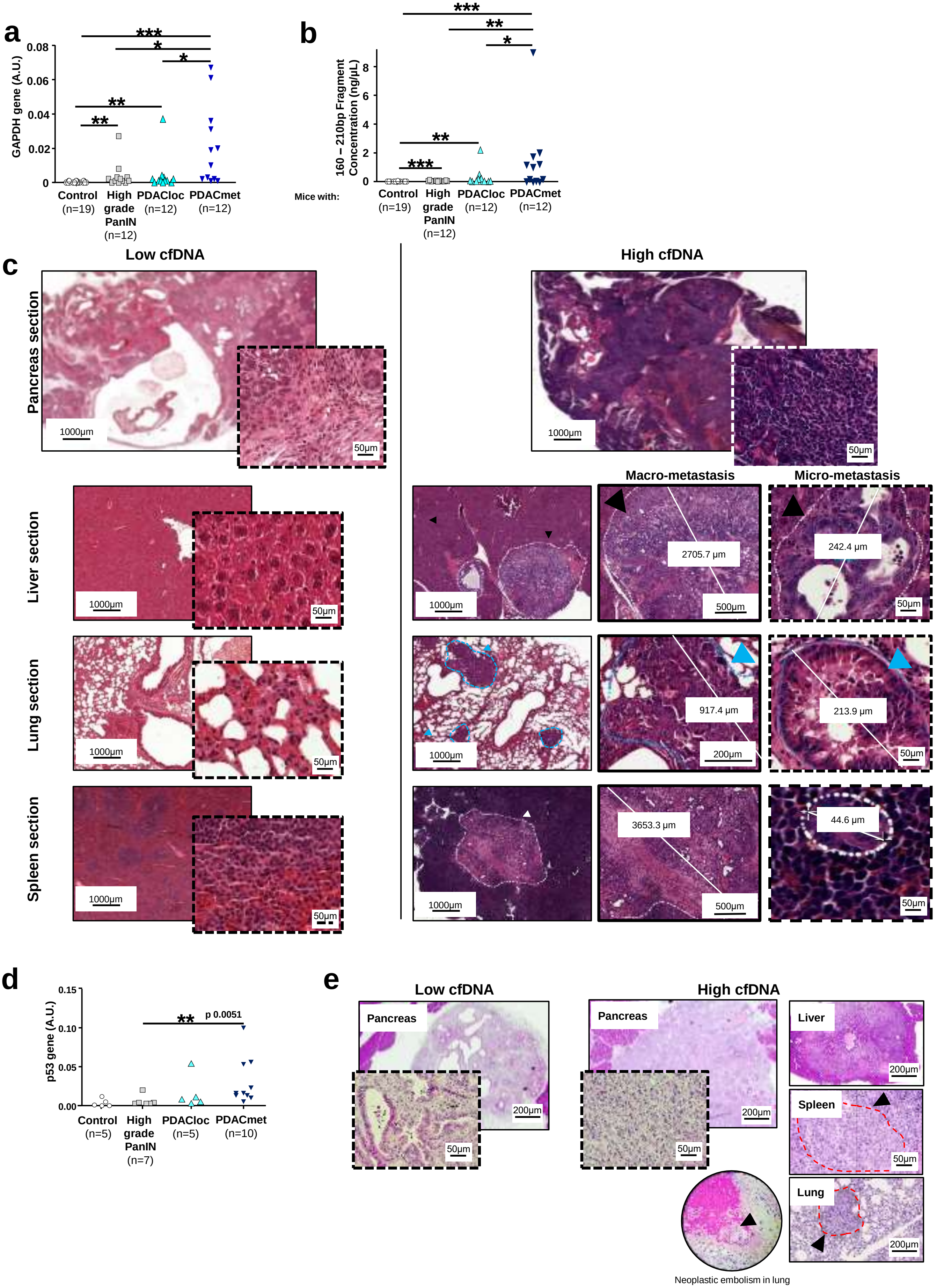


#### Supplementary Figure 10

### S13

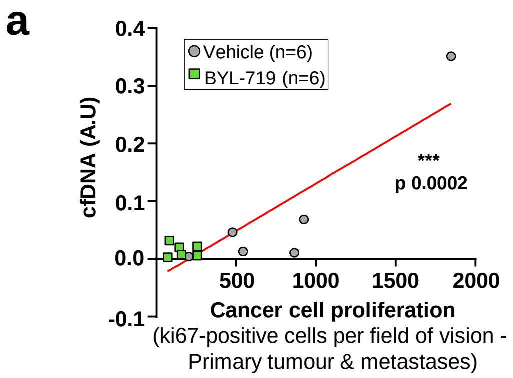


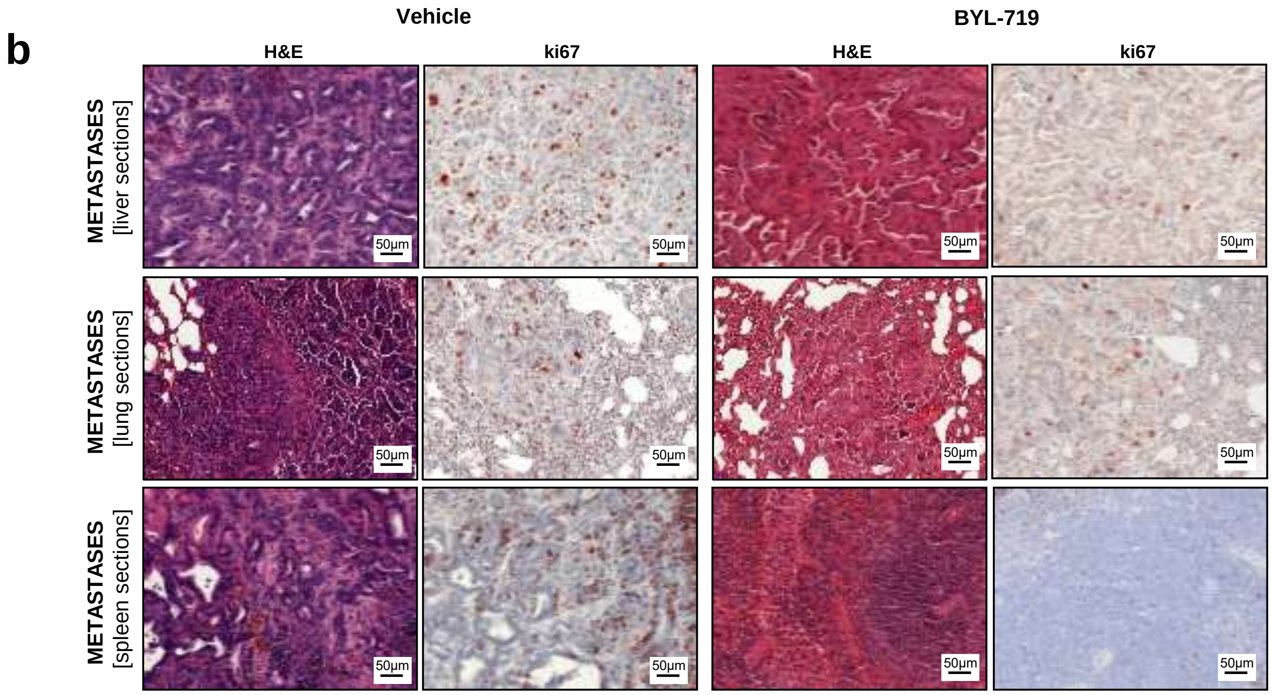


#### Supplementary Figure 13

### S15

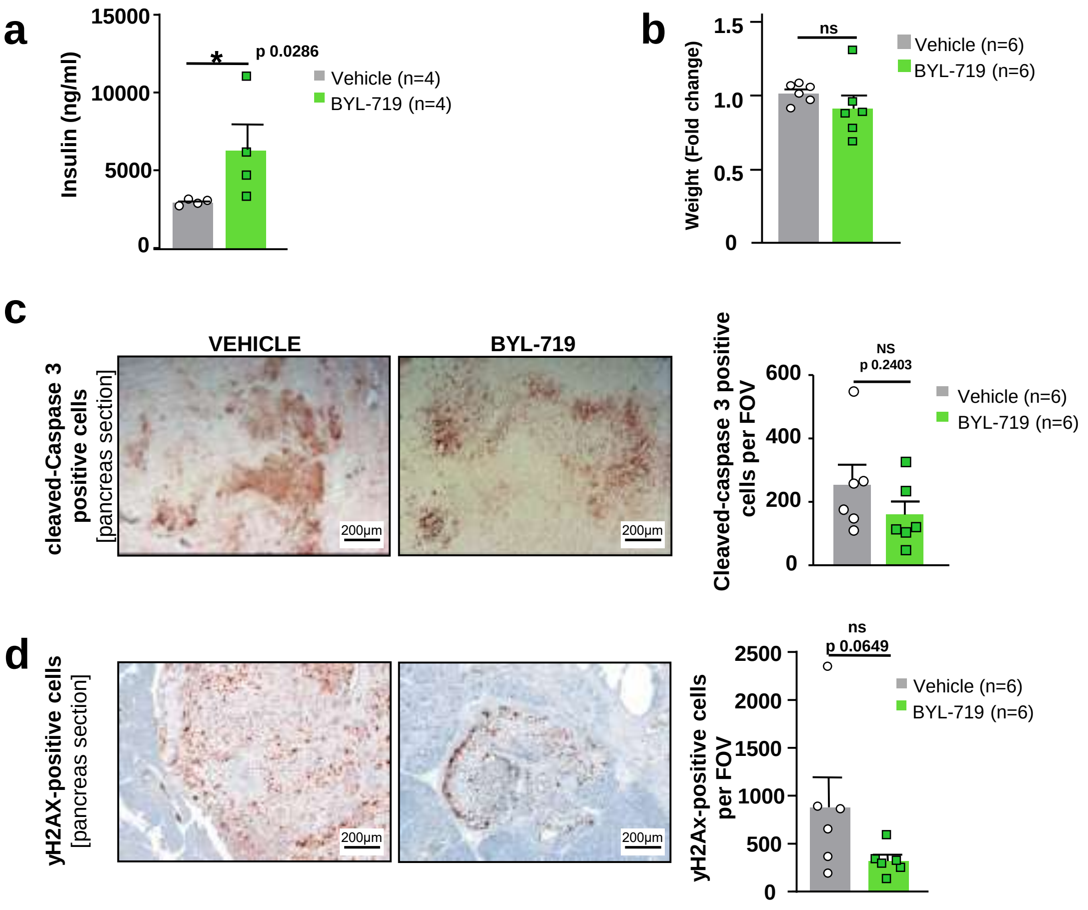


**Supplementary Figure 15**

### S16

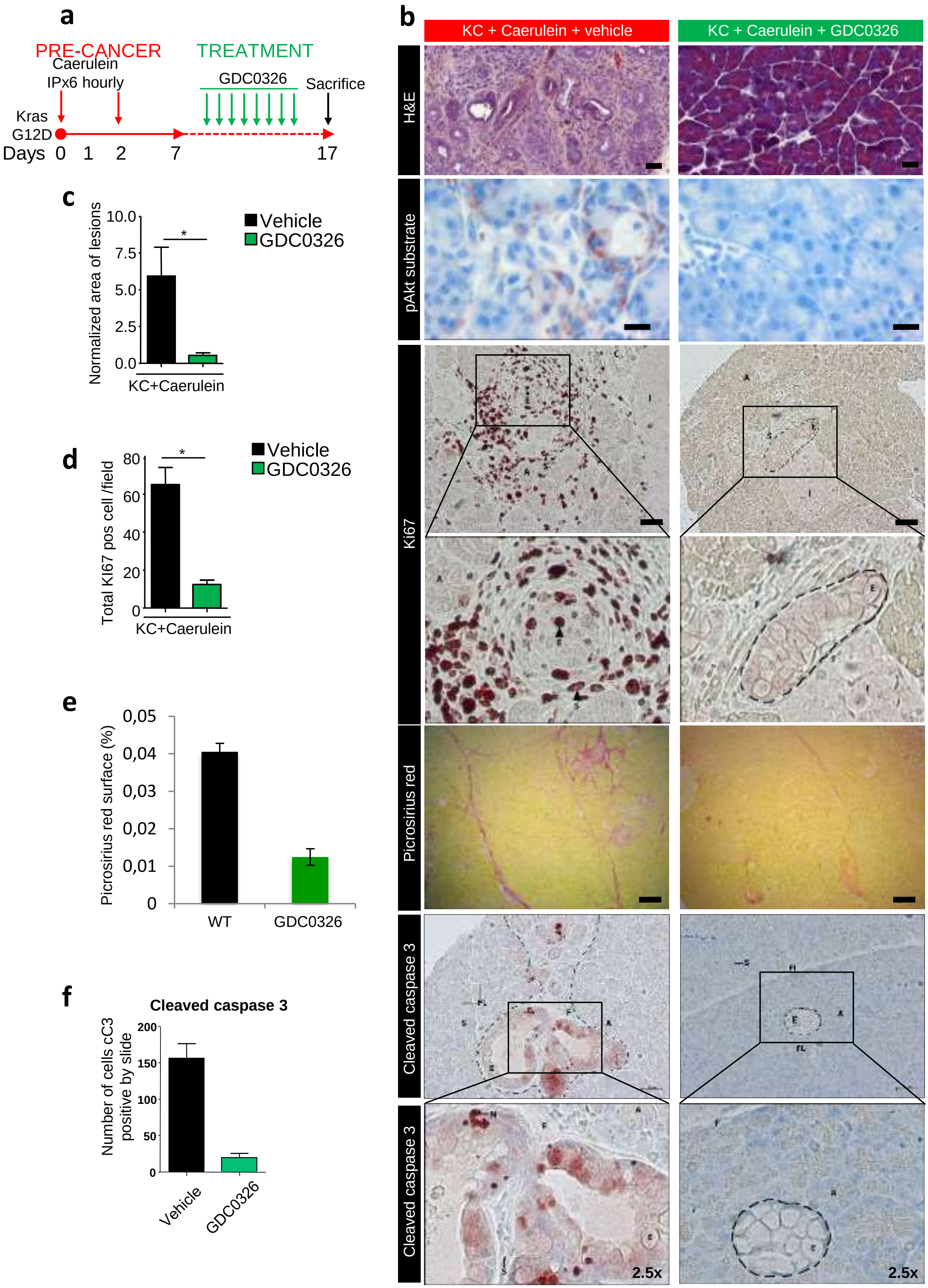


**Supplementary Figure 16**

### S17

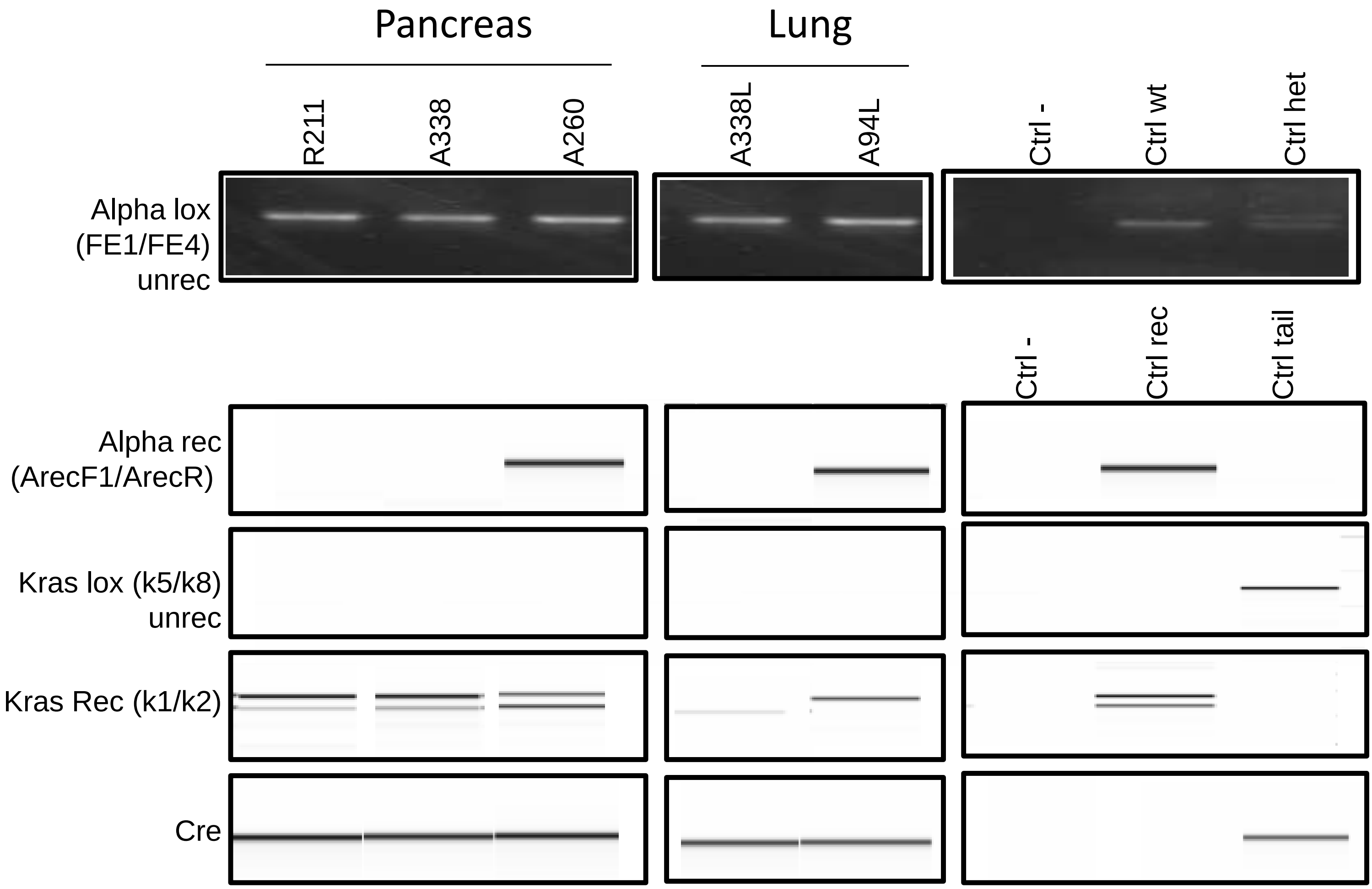


**Supplementary Figure 17**

### S18

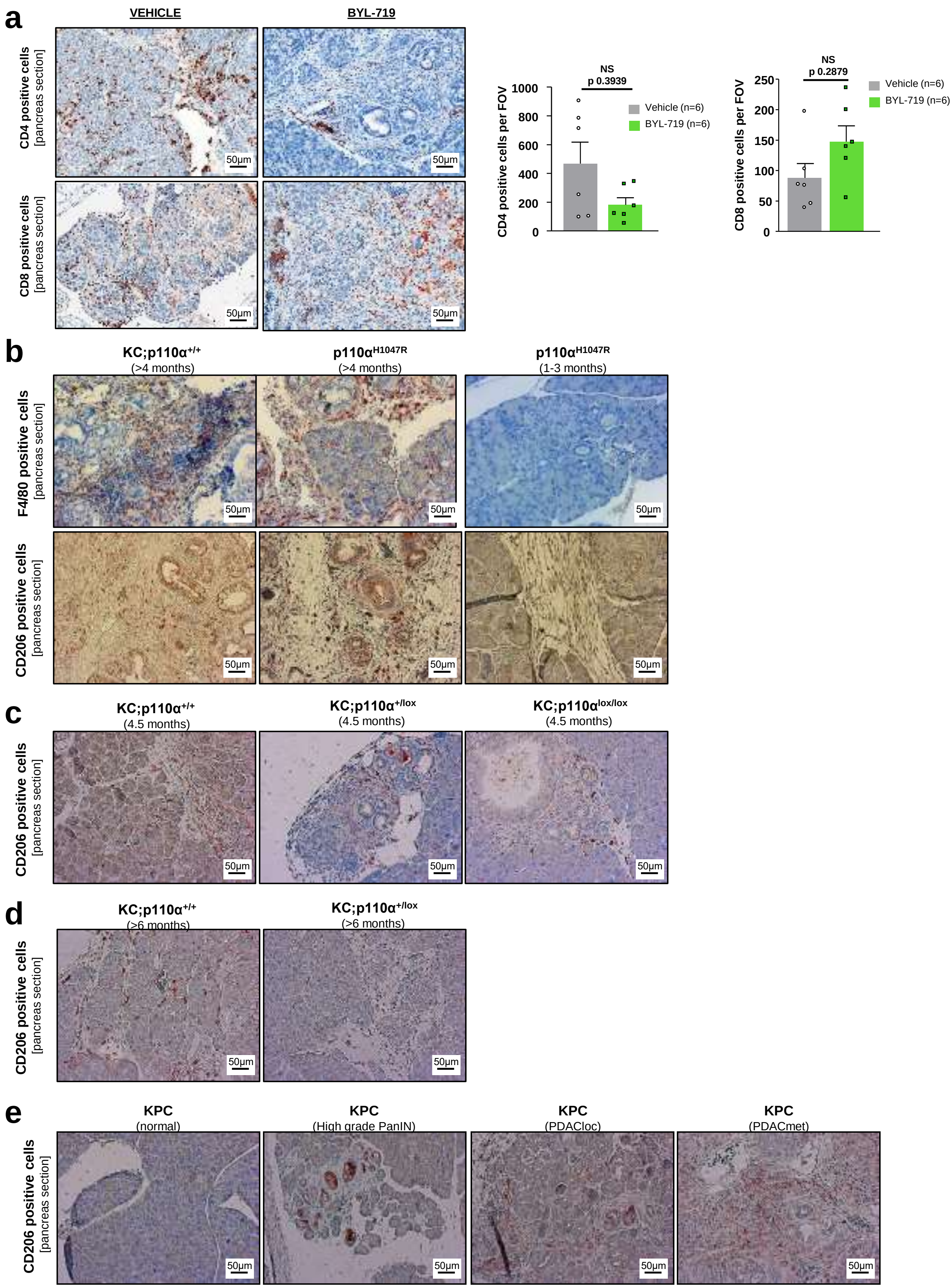


#### Supplementary Figure 18
