## Supplementary material for "Pancreatic Cancer Intrinsic PI3Kα Activity accelerates Metastasis and rewires Macrophage Component": S6

**a**

| | Cytotoxicity - Growth GI30 (µM) | | | | | | | | | | | |
|---|---|---|---|---|---|---|---|---|---|---|---|---|
| | Cell line | A66 | BYL-719 | TGX-221 | TGX-155 | TGX-115 | AZD8186 | AS252424 | LY294002 | BKM120 | GDC0941 | PI-103 |
| | | α | | β | | β/δ | | γ | pan | | | |
| Pancreas | R211 | NR | 5.804 | NR | NR | NR | NR | NR | 2.097 | 1.307 | 1.699 | 1.878 |
| | PDAC8661 | NR | 1.867 | NR | NR | NR | NR | NR | 9.575 | 1.022 | 1.127 | 6.389 |
| | DT4994 | NR | 6.082 | NR | NR | NR | NR | NR | 9.096 | 2.370 | 2.043 | 8.421 |
| | PANC-1 | NR | NR | NR | NR | NR | NR | NR | 4.15 | 1.638 | 1.654 | 0.818 |
| | 10593 | 2.017 | 0.8107 | - | - | - | - | NR | - | 0.615 | 0.410 | - |
| | 10158 | 1.182 | 0.6293 | - | - | - | - | NR | - | 0.545 | 0.312 | - |
| | R6344 | 1.443 | 1.311 | 6.663 | NR | NR | 2.662 | NR | 3.556 | 0.944 | 0.501 | 6.288 |
| | R6430 | 1.487 | 1.955 | NR | NR | NR | 2.617 | NR | 2.471 | 0.723 | 0.359 | 0.646 |
| | R6065 | 2.787 | 1.928 | NR | NR | NR | 2.887 | NR | 8.330 | 0.701 | 0.331 | 2.528 |
| | R6141 | 7.519 | 7.608 | NR | NR | NR | 5.753 | NR | 8.477 | 1.845 | 0.782 | 2.965 |
| Other organs | PC3 | NR | 8.671 | 2.005 | NR | NR | 0.3351 | NR | 4.203 | 0.660 | 0.217 | 0.058 |
| | MDA-MB-231 | NR | NR | 3.832 | NR | NR | NR | NR | 3.163 | 0.794 | 1.296 | 0.217 |
| | MDA-MB-468 | NR | NR | 3.800 | NR | 7.790 | 1.365 | NR | 8.549 | 0.989 | 0.513 | 0.206 |
| | NOMO-1 | NR | 4.291 | NR | NR | NR | NR | 8.741 | 7.341 | 1.096 | 0.319 | 0.727 |
| | HL-60 | NR | NR | NR | NR | NR | NR | 8.326 | NR | 0.583 | 0.629 | 1.284 |

| | Cytotoxicity - Growth GI30 (µM) | | | | | | | | | | | |
|---|---|---|---|---|---|---|---|---|---|---|---|---|
| | Cell line | A66 | BYL-719 | TGX-221 | TGX-155 | TGX-115 | AZD8186 | AS252424 | LY294002 | BKM120 | GDC0941 | PI-103 |
| | | α | | β | | β/δ | | γ | pan | | | |
| Pancreas | HPNE hTERT | 3.644 | 4.141 | NR | NR | NR | NR | NR | 8.801 | 1.174 | 0.541 | 1.366 |

NR = not reached
- = non-available

**b**

| | Migration - Inhibitory IC30 (µM) | | | | | | | | | | | |
|---|---|---|---|---|---|---|---|---|---|---|---|---|
| | Cell line | A66 | BYL-719 | TGX-221 | TGX-155 | TGX-115 | AZD8186 | AS252424 | LY294002 | BKM120 | GDC0941 | PI-103 |
| | | α | | β | | β/δ | | γ | pan | | | |
| Pancreas | R211 | 0.045 | 0.044 | NR | NR | NR | NR | NR | NR | NR | 0.035 | 0.151 |
| | PDAC8661 | 0.814 | 0.177 | NR | NR | NR | NR | NR | NR | NR | 0.053 | 0.137 |
| | PANC-1 | 0.133 | 0.148 | 0.8217 | NR | NR | NR | NR | NR | NR | 0.105 | 0.187 |
| | 10593 | 0.854 | 0.171 | - | - | - | - | NR | - | 0.511 | 0.033 | - |
| | 10158 | 0.694 | 0.417 | - | - | - | - | NR | - | 0.918 | 0.038 | - |
| | R6344 | NR | 0.375 | NR | NR | NR | NR | NR | NR | NR | 0.416 | 0.463 |
| | R6430 | 0.986 | 0.132 | NR | NR | NR | NR | NR | NR | 0.536 | 0.163 | 0.198 |
| | R6065 | 0.421 | 0.076 | 0.028 | 0.846 | NR | 0.026 | NR | NR | 0.109 | 0.026 | 0.075 |
| | R6141 | NR | 0.198 | NR | NR | 0.941 | 0.033 | NR | NR | 0.604 | 0.114 | 0.048 |
| Other organs | PC3 | NR | NR | 0.048 | 0.548 | 0.014 | 0.028 | NR | NR | 0.743 | 0.056 | 0.064 |
| | MDA-MB-231 | NR | NR | 0.861 | NR | NR | 0.013 | NR | NR | NR | 0.101 | 0.030 |
| | MDA-MB-468 | NR | NR | 0.252 | NR | 0.963 | 0.012 | NR | NR | NR | 0.095 | 0.021 |

| | Migration - Inhibitory IC30 (µM) | | | | | | | | | | | |
|---|---|---|---|---|---|---|---|---|---|---|---|---|
| | Cell line | A66 | BYL-719 | TGX-221 | TGX-155 | TGX-115 | AZD8186 | AS252424 | LY294002 | BKM120 | GDC0941 | PI-103 |
| | | α | | β | | β/δ | | γ | pan | | | |
| Pancreas | HPNE hTERT | 0.989 | NR | NR | NR | NR | 0.921 | NR | NR | NR | 0.244 | 0.148 |

NR = not reached
- = non-available

### Supplementary Figure 6
