## Supplementary material for "Pancreatic Cancer Intrinsic PI3Kα Activity accelerates Metastasis and rewires Macrophage Component": S9

**a**

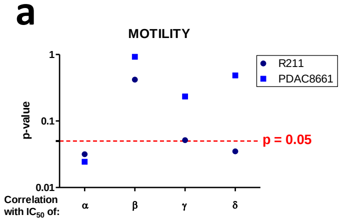


**b**

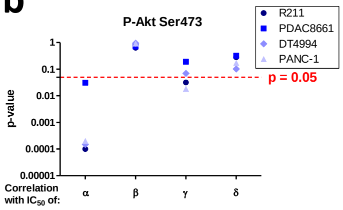


**c**

| | Migration correlation - R squared | | | | |
|---|---|---|---|---|---|
| | | PI3Kα | PI3Kβ | PI3Kγ | PI3Kδ |
| Pancreas | **R211** | 0.6707 | 0.01982 | 0.2436 | 0.1686 |
| | **PDAC8661** | 0.7075 | 0.0005708 | 0.3281 | 0.2799 |
| | **PANC-1** | 0.5336 | 0.0397 | 0.1811 | 0.3739 |
| | **10593** | 0.8969 | 0.6883 | 0.04761 | 0.8156 |
| | **10158** | 0.792 | 0.3554 | 0.354 | 0.5543 |
| | **R6344** | 0.388 | 0.06159 | 0.1963 | 0.3665 |
| | **R6430** | 0.6669 | 0.001815 | 0.3333 | 0.2532 |
| | **R6065** | 0.6097 | 0.2529 | 0.2354 | 0.6925 |
| | **R6141** | 0.5217 | 0.1696 | 0.2579 | 0.6677 |
| Other organs | **PC3** | 0.06767 | 0.8464 | 0.07215 | 0.7513 |
| | **MDA-MB-231** | 0.1328 | 0.1938 | 0.1641 | 0.3925 |
| | **MDA-MB-468** | 0.05677 | 0.4916 | 0.05836 | 0.5726 |

**d**

| | Cytotoxicity correlation - R squared | | | | |
|---|---|---|---|---|---|
| | | PI3Kα | PI3Kβ | PI3Kγ | PI3Kδ |
| Pancreas | **R211** | 0.3917 | 0.01859 | 0.2506 | 0.2026 |
| | **PDAC8661** | 0.5084 | 0.01331 | 0.3061 | 0.2586 |
| | **DT4994** | 0.5554 | 0.02876 | 0.3779 | 0.2945 |
| | **PANC-1** | 0.3131 | 0.1128 | 0.2374 | 0.3323 |
| | **10593** | 0.7159 | 0.6617 | 0.113 | 0.7343 |
| | **10158** | 0.7154 | 0.527 | 0.2357 | 0.6636 |
| | **R6344** | 0.7754 | 0.01008 | 0.3046 | 0.2667 |
| | **R6430** | 0.6886 | 0.007813 | 0.238 | 0.2871 |
| | **R6065** | 0.7842 | 0.005775 | 0.3083 | 0.2899 |
| | **R6141** | 0.6351 | 0.0555 | 0.3203 | 0.4164 |
| Other organs | **PC3** | 0.3739 | 0.3731 | 0.2671 | 0.6739 |
| | **MDA-MB-231** | 0.2255 | 0.07722 | 0.2499 | 0.2245 |
| | **MDA-MB-468** | 0.2327 | 0.2938 | 0.2018 | 0.4658 |
| | **NOMO-1** | 0.5756 | 0.006474 | 0.5803 | 0.169 |
| | **HL-60** | 0.4795 | 0.001741 | 0.7097 | 0.09689 |

**e**

| | Motility correlation - R squared | | | | |
|---|---|---|---|---|---|
| | | PI3Kα | PI3Kβ | PI3Kγ | PI3Kδ |
| Pancreas | **R211** | 0.8291 | 0.2247 | 0.7671 | 0.8178 |
| | **PDAC8661** | 0.8556 | 0.003624 | 0.4237 | 0.174 |

**f**

| | p-Akt Ser473 correlation - R squared | | | | |
|---|---|---|---|---|---|
| | | PI3Kα | PI3Kβ | PI3Kγ | PI3Kδ |
| Pancreas | **R211** | 0.8288 | 0.02511 | 0.4175 | 0.1272 |
| | **PDAC8661** | 0.4189 | 0.01634 | 0.1822 | 0.109 |
| | **DT4994** | 0.8141 | 0.0004657 | 0.3192 | 0.2686 |
| | **PANC-1** | 0.8671 | 0.001017 | 0.4797 | 0.1964 |

### Supplementary Figure 9
