## Supplementary material for "Pancreatic Cancer Intrinsic PI3Kα Activity accelerates Metastasis and rewires Macrophage Component": S11

a

| Enrolment criteria for KPC mice | |
|---|---|
| Detected tumour by US imaging | |
| Tumour location | Head or tail |
| Increased cfDNA* | |
| Weight | > 22g |
| Sex | Male or female |

b

| Mouse ID | Sex | Age (weeks) | Initial weight (g) | cfDNA (Fold increase) | Tumour location | Treatment |
|---|---|---|---|---|---|---|
| 637 | M | 35 | 25.6 | 2.333 | Head | BYL-719 |
| 715 | M | 32 | 27.3 | 4.000 | Head | BYL-719 |
| 773 | F | 29 | 25.2 | 1.000 | Head | BYL-719 |
| 785 | M | 30 | 32.5 | 0.500 | Tail | BYL-719 |
| 871 | M | 24 | 36.3 | 0.750 | Tail | BYL-719 |
| 897 | M | 21 | 36.1 | 5.750 | Head | BYL-719 |
| 790 | M | 32 | 28 | 1.500 | Tail | Vehicle |
| 795 | F | 33 | 24.7 | 1.667 | Head | Vehicle |
| 860 | M | 23 | 30.4 | 0.222 | Head | Vehicle |
| 894 | M | 25 | 32 | 3.500 | Tail | Vehicle |
| 939 | F | 18 | 22.3 | 2.167 | Head | Vehicle |
| 977 | M | 20 | 30.9 | 6.333 | Tail | Vehicle |

### Supplementary Figure 11
