## Supplementary material for "Pancreatic Cancer Intrinsic PI3Kα Activity accelerates Metastasis and rewires Macrophage Component": S12

**a**

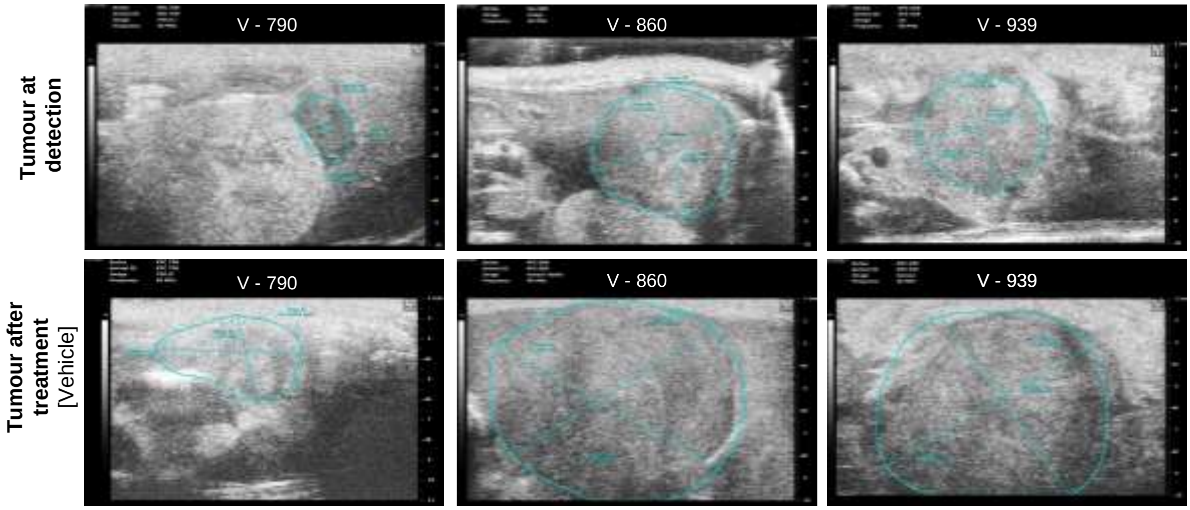


**b**

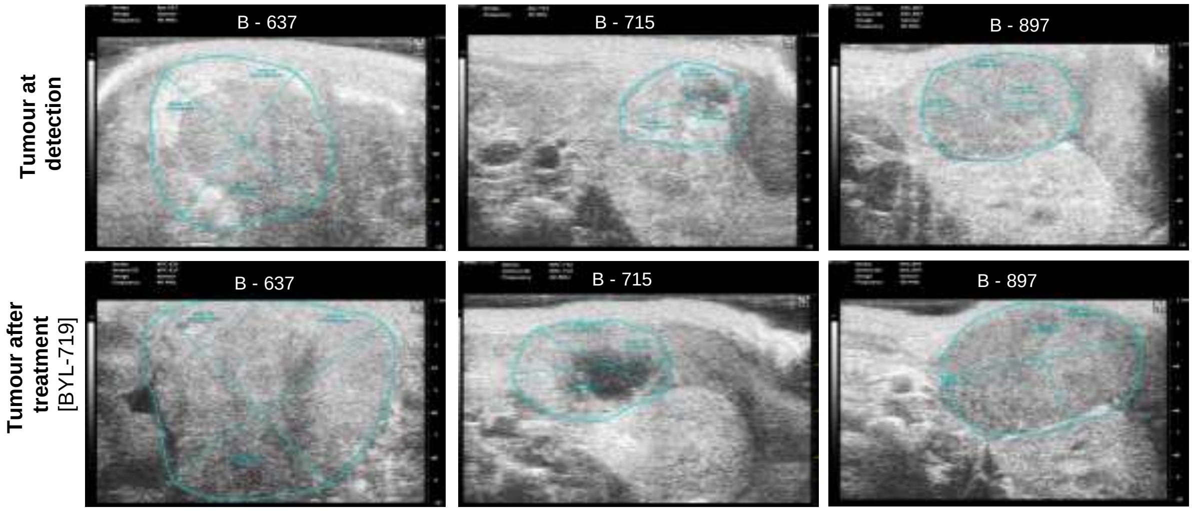


**c**

| Mouse ID | Initial Tumour volume ($mm^3$) | Final Tumour volume ($mm^3$) | Tumour fold change | Treatment |
|---|---|---|---|---|
| **637** | 435.523 | 656.492 | 1.507 | BYL-719 |
| **715** | 62.045 | 97.522 | 1.572 | BYL-719 |
| **773** | 156.436 | 555.745 | 3.553 | BYL-719 |
| **785** | 256.423 | 403.214 | 1.572 | BYL-719 |
| **871** | 288.199 | 357.401 | 1.240 | BYL-719 |
| **897** | 181.604 | 265.693 | 1.463 | BYL-719 |
| **790** | 14.671 | 154.878 | 10.557 | Vehicle |
| **795** | 450.402 | 821.260 | 1.823 | Vehicle |
| **860** | 465.018 | 1045.508 | 2.248 | Vehicle |
| **894** | 179.036 | 458.345 | 2.560 | Vehicle |
| **939** | 93.553 | 620.745 | 6.635 | Vehicle |
| **977** | 435.572 | 1041.951 | 2.392 | Vehicle |

### Supplementary Figure 12
