## Supplementary material for "Pancreatic Cancer Intrinsic PI3Kα Activity accelerates Metastasis and rewires Macrophage Component": S14

**a**

| Mouse ID | Treatment | Tumour location | Primary tumour | Histological Grade | Regional lymph nodes | Distant Metastases | Metastasic sites | Stage |
|---|---|---|---|---|---|---|---|---|
| B-637 | BYL-719 | Head | T4 | G3 | N1 | M1 | Liver, lung, peritoneum, | IV |
| B-715 | BYL-719 | Head | T2 | G2 | N0 | M0 | N/A | IB |
| B-773 | BYL-719 | Head | T4 | G3 | N1 | M1 | Liver, peritoneum, ascites | IV |
| B-785 | BYL-719 | Tail | T4 | G3 | N0 | M0 | N/A | III |
| B-871 | BYL-719 | Tail | T3 | G3 | N0 | M0 | N/A | III |
| B-897 | BYL-719 | Head | T4 | G3 | N1 | M1 | Liver | IV |
| V-790 | Vehicle | Tail | T2 | G4 | N0 | M0 | N/A | IB |
| V-795 | Vehicle | Head | T4 | G3 | N1 | M1 | Liver, lung, spleen, peritoneum, ascites | IV |
| V-860 | Vehicle | Head | T4 | G3 | N1 | M1 | Liver, spleen, peritoneum, ascites | IV |
| V-894 | Vehicle | Tail | T4 | G4 | N1 | M1 | Liver, spleen, peritoneum, ascites | IV |
| V-939 | Vehicle | Head | T4 | G4 | N1 | M1 | Liver, lung, peritoneum, ascites | IV |
| V-977 | Vehicle | Tail | T4 | G4 | NX | M1 | Liver, peritoneum | IV |

***Primary Tumour (T)***
TX Primary tumour cannot be assessed
T0 No evidence of primary tumour
Tis Carcinoma in situ* includes PanIN III
T1 Tumour limited to the pancreas, less than 2mm in biggest dimension
T2 Tumour limited to the pancreas, more than 2mm in biggest dimension
T3 Tumour extends beyond the pancreas but without involvement of the celiac axis or main arteries
T4 Tumour involves celiac axis or main arteries (unresectable primary tumour)

***Regional Lymph Nodes (N)***
NX Regional lymph nodes cannot be assessed
N0 No regional lymph node metastasis
N1 Regional lymph node metastasis

***Distant Metastases (M)***
M0 No distant metastases
M1 Distant metastases

***Histological grade (G)***
GX cannot be assessed
G1 well differentiated
G2 moderately differentiated
G3 poorly differentiated
G4 undifferentiated

**b**

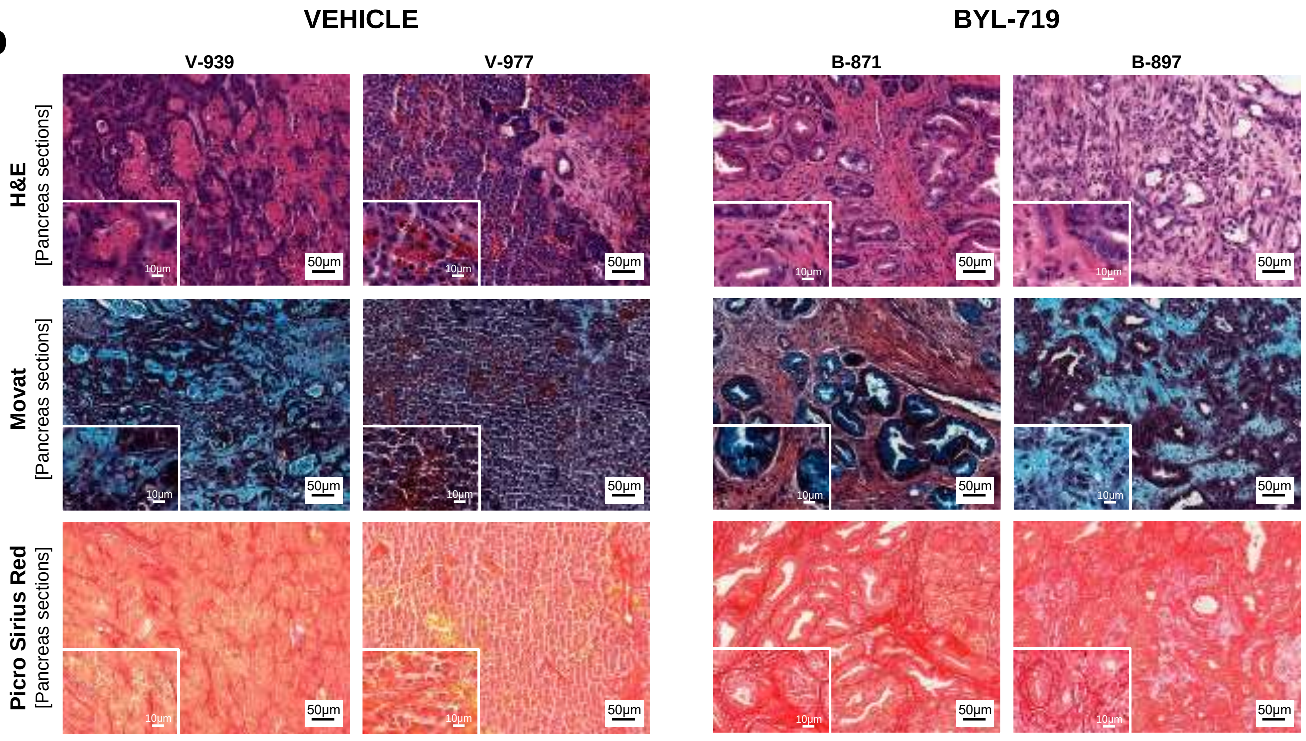


### Supplementary Figure 14
